# Barking Up the Right Tree: Immune Checkpoint Signatures of Human and Dog Cancers

**DOI:** 10.1101/2024.06.26.600825

**Authors:** Mikolaj Kocikowski, Marcos Yébenes Mayordomo, Javier Alfaro, Maciej Parys

## Abstract

In the quest for improved therapeutics targeting immune checkpoints (ICs), we turn to spontaneously developing dog (canine) cancers, which are unique models that genetically and clinically mirror human equivalents. Despite its potential, canine cancer immunology remains largely unexplored. Here, we examine the RNA-seq-based expression of 44 ICs across 14 canine cancer types and an extensive human dataset. We unveil diverse canine IC expression patterns and unique human IC signatures that reflect the histological type and primary site of cancer. We uncover a striking similarity between canine brain cancers, osteosarcoma, and their human counterparts, identifying them as prospective immunotherapy models. Four ICs—CD160, A2AR, NKG2A, and OX40—are key to the differences observed between species. Moreover, individual patient IC signatures exhibit varying alignment with their respective cancer types, a finding with profound implications for personalized human therapy. This exploration illuminates new aspects of canine and human cancer immunology, setting the stage for discoveries at their crossroads.

## Introduction

Immune checkpoints (ICs) are signaling pathways that play a key role in regulating immune responses by modulating the activity of immune cells. Disruption of these pathways has shown promise in cancer treatment, leading to the development of immune checkpoint blockers (ICB; or inhibitors - ICI) as a therapeutic strategy. Antibody-based ICB therapeutics induce remarkable remissions in some cancer cases^1, 2^. While their efficacy is limited to a subset of patients, adverse effects affect more than 60% of treated individuals and may include rapid cancer progression^3–5^. The extent of unforeseen adverse effects necessitates a more robust approach to immunotherapy development amid the surge in targeting underexplored ICs and their combinations^5–8^. Novel cancer models, such as canine (dog) cancers, hold potential to address this challenge.

Unlike conventional chemotherapy, ICB immunotherapy modulates a complex signaling network to reactivate patient’s immune cells so that they can recognize and combat immune-edited cancer^9^. The curative and toxic outcomes of this intricate process cannot be thoroughly evaluated in rodent models based on xenografts injected into immunocompromised animals or syngeneic, artificially induced tumors. In contrast, canine cancers develop organically under the surveillance of a complete immune system, reflecting the complexities of naturally occurring human cancers. Dogs develop a spectrum of cancers, some common and others rare in humans. For the rare ones, such as pediatric osteosarcoma^10, 11^, canine cancers enable otherwise improbable, high-powered studies. Several canine cancers mirror their human counterparts at the histological, clinical and genetic level^12^. Consequently, they rise as prospective research models^13, 14^. More than 40 immune checkpoints have been identified, but the presence of these potential immunotherapy targets in canine cancers remains mostly unknown^15^. The feasibility of canine immunotherapy hinges upon comprehensive characterization of the canine IC landscape. Only canine cancers that show immune checkpoint patterns similar to those found in human cancers can serve as valuable models for immunotherapy studies.

In contrast to that in dogs, much is known about the abundance of individual ICs in most human cancers^16^. Expression patterns encompassing multiple ICs have been linked to treatment responses and prognosis in several cancer types, which supports the clinical value of considering multi-IC patterns^17–20^. However, comprehensive IC signatures have not been investigated. Clinically important yet unanswered questions include the influence of cancer histological type on IC abundance patterns, the existence of immunologically related cancer clusters, and the similarity of signatures among patients with the same cancer type.

In this study, we pursued three objectives: first, to assess the abundance of immune checkpoints across canine cancer types; second, to compare these IC profiles with those found in human cancers; and third, to analyze the IC patterns within a species and between individual patients. We began by integrating gene expression data encompassing 14 canine and 27 human cancer types. We detected the expression of all ICs in canine cancers, albeit the levels varied greatly amongst the types. Subsequently, we generated human and canine cancer type-specific IC signatures. We compared these signatures and found similarities between cancers of both species. These similarities were particularly striking in the case of brain cancers. We also observed exceptional similarity between canine osteosarcoma and human sarcoma. In hierarchical clustering, most cancer signatures aggregated by species, followed by histological type and primary site. A2AR and possibly six other IC genes drove the species divide. When examining individual patient IC signatures, we found that patients with the same cancer type tended to cluster together, although diverse signatures were observed in certain cancer types, such as prostate carcinoma. Furthermore, we discovered unexpected similarities between human chronic lymphocytic leukemia cases and canine malignancies at large. We mapped the immune checkpoint landscape of canine and human cancers, revealing shared expression patterns and possible immunotherapy targets. Our findings underscore the untapped potential of cancer research in dogs and further the understanding of IC signatures, a step towards advancing targeted immunotherapy.

## Materials and Methods

### Acquisition of Canine Gene Expression Data

Openly available canine RNA sequencing data were utilized for this analysis. A comprehensive literature search was conducted on PubMed on 2022-05-15, employing the following search query: ‘((rnaseq) OR (rna-seq) OR (rna sequencing) OR (transcriptomics)) AND ((dog) OR (canine) OR (canis)) AND cancer’. Of the 413 identified articles, 15 studies representing 8 cancer types met the inclusion criteria outlined below:

1. Pertinence to canine cancer.
2. Inclusion of original data.
3. Data source: data originated from tumor tissues as opposed to cell cultures, single cells or exosomes; cell lines are notoriously different from their tumors of origin, and data for single cells or extracellular vesicles are not comparable to those for bulk tumors.
4. Accessibility: RNAseq data were declared openly available, linked to and accessible.
5. Data format: raw data were provided as FASTQ, SRA or BAM (complete) files so that the raw reads from all the datasets could be analyzed together consistently.
6. Adequate description: the samples and methods were sufficiently described, ensuring the used protocols make the data reliable and comparable.
7. Sample preservation method: samples were preserved by snap-freezing or RNAlater infusion and freezing, which is known to maintain RNA quality better than the FFPE method.
8. mRNA enrichment: Samples were mRNA-enriched through poly-A selection or rRNA depletion, as total RNA and mRNA sequencing would not be fully comparable.
9. Sequencing method: Paired-end (PE) sequencing was conducted, providing more accurate and reliable data by using contemporary sequencing technology.
10. Treatment status: None of the animals appeared to have received chemotherapy or radiation therapy before tissue sampling, as this could alter the transcriptome of the tumors, including IC expression. This study focused on treatment-naive tumors.

If multiple studies of the same cancer type passed the inclusion criteria, the one with the maximum sample size was included. Additionally, pulmonary neoplasm and oral melanoma datasets unavailable elsewhere were obtained from the ICDC canine commons. We strived to have no less than 10 samples per cancer type where possible; hence, an additional T-cell lymphoma dataset was downloaded from the ICDC. All the ICDC data met our inclusion criteria. In the originally chosen HSA study, 15 of 23 samples were rejected because of very short (length: 25) reads that were incongruent with the other samples (length: 38) and not compatible with the quantification settings. To obtain a sufficient number of samples, we used data from an additional publication on hemangiosarcoma (of the 51 samples found in the SRA database, 6 appeared to be derived from cell lines and were excluded, leaving 45 samples, 2 fewer than those described in the paper).

### Exceptions

We made the following exceptions to obtain insight into cancer types unavailable otherwise: OSCC (FFPE samples), ameloblastoma (SE sequencing), meningioma (SE + FFPE), and prostate cancer (total RNA library). In the case of insulinoma, the type of sequenced RNA library was unknown, the reads were pre-trimmed with an unknown method, and the shared bam files were based on an older canine reference genome (CanFam3.1). Hence, insulinoma analysis must be considered purely exploratory. One of the 12 ameloblastoma samples was not downloadable at the time of the analysis. In the BCL study, 4 of 16 patients had either T-cell lymphoma or undefined lymphoma. In the glioma study, of the 83 samples, 38 were FFPE, and of the 45 frozen samples, 42 were available for download. The samples that did not satisfy the inclusion criteria were excluded. A summary of the cancer types, their abbreviations, data sources and articles of origin can be found in **Table S5**.

### Acquisition of Human Gene Expression Data

The quantified expression of the IC genes of interest was sourced from the PanCancer Analysis of Whole Genomes (PCAWG) dataset and obtained from the European Bioinformatics Institute (EBI) Expression Atlas. The dataset encompassed 1200 samples from 27 human cancer types. Detailed descriptions of the data acquisition, ethical considerations, and sequencing techniques can be found in their accompanying publication^21^.

### Processing of Canine RNA Sequencing Data

The raw canine RNA-seq data were preprocessed from BAM or SRA files into fastq format, subjected to quality control with FastQC v0.11.6 (bioinformatics.bbsrc.ac.uk/projects/fastqc) and MultiQC^22^ v1.12, and quantified without trimming on a PLGrid Prometheus computational cluster. Reads were quantified with Kallisto^23^ v0.45.0 using a Kallisto-generated index based on an Ensembl canine reference transcriptome release 106 (Canis_lupus_familiarisboxer.Dog10K_Boxer_Tasha.cdna.all.fa file version). The parameters for the PE data were default (K-mer length of 31). The setting for SE samples was “-l 200 -s 20”, as bioanalyzer reports for sequencing libraries were not available. The resulting abundance data were imported to R/RStudio using the following libraries: GenomicFeatures^24^, tximport, and tximportData^25^. Technical replicates were collapsed, the entire dataset was normalized with DESeq2^26^, and the results were visualized using ggplot2^27^ combined with the *Viridis* colourblind-friendly colour map^28^. DESeq2 performs normalization in regard to the size and composition of sequencing libraries created in each experiment, which allows for a comparison of different sample groups (cancer types in our case). When combined with tximport, it also normalizes the abundance data in regard to the average transcript length of each gene. This in turn allows for comparison of expression between different genes. For all canine (single-species) analyses, DESeq-2-normalized counts were used as the most appropriate for combining sequencing data from different protocols and tissues. The relationship between gene expression and particular dog breeds was not assessed due to the inconsistent quality of the provided sample metadata.

### Integration of Human and Canine Data

To compare cross-species datasets, the dataset representing the abundance of IC expression in canine cancers was transformed into transcripts per million (TPM) units. This version of the dataset will be referred to as “TPM data” throughout the remainder of this section. This transformation was carried out to ensure compatibility between the canine data and the PCAWG human dataset, which was already quantified in TPM. Canine and human datasets were merged and standardized (scaled and centered) using the ‘scale’ function in R. A standardized dataset was used for subsequent analyses unless otherwise stated.

### Selection of Immune Checkpoints

Research on immune checkpoints is dynamically evolving, and new ICs emerge. A list of IC receptors and ligands, including well-known (PD-1) and emerging (SLAMF7) ones, was generated based on the literature^29^. ICs that were available in the canine reference transcriptome of choice were selected for the study (**Tab. S6**).

### Conservation of the IC Amino Acid Sequence

The identity [%] and coverage [%] were calculated by BLASTP alignment of canine and human amino acid sequences of the top canonical protein-coding Ensembl transcripts. The conservation score was calculated as identity*coverage/100. One caveat of this approach is that it does not prioritize the extracellular, exposed domains (potential antibody epitopes). The values are included in **Table S6.**

### Selection of Cell-Marker Genes

Our list of immune cell markers was inspired by the NanoString panels^30^. The list was adapted based on the reference transcriptome and gene specificity in the Human Protein Atlas^31^. The TNFRSF17 gene was rejected because it was absent in the quantification results, and the CPA3 gene was rejected because of misleading results (see Supplementary Methods).

### Immune Infiltrate Composition

For each of the chosen marker genes, a mean value was calculated based on its means in all cancer types. Subsequently, expression levels for individual cancer types were normalized by dividing them by this mean. This normalization approach was adopted to ensure equal contributions from each marker to the overall analysis, independent of their inherent abundance. In the absence of precise information regarding canine cell markers, this strategy was deemed most appropriate. For each cell type category, means of normalized values were computed for marker genes. These averaged values were then visualized.

### Immune Inhibition Score

The immune inhibition score was calculated in a manner analogous to how the immune infiltrate composition was estimated. Immune checkpoints (ICs) were categorized as ‘inhibitory’, ‘stimulatory’, or ‘other’. For each IC, the mean expression value was calculated across cancer types. The means for individual cancer types were subsequently divided by this mean to account for the wide variations in IC expression. The data were then averaged within each category. For each cancer type, the derived inhibitory score was divided by the stimulatory score, resulting in a singular estimate that represented the immune environment character for that particular cancer.

### Hierarchical Clustering (HC)

Hierarchical clustering was conducted in R based on standardized cross-species TPM data. For each cancer type, a distinct vector was created to encapsulate the median expression of all ICs. Those vectors were compared in a distance matrix created with the ‘amap::Dist’ package employing Pearson’s correlation as the distance metric. The resulting matrix was subjected to hierarchical clustering using the ‘hclust’ function with the ‘ward.D2’ Ward linkage method. The clustered output was visualized as a dendrogram with the aid of the ‘factoextra’ package in R.

### Differential Transcript Abundance Between Clusters

An exploratory analysis of differential transcript abundance between cancer type clusters was performed in R based on the TPM data.

### Functional Network Analysis of Immune Checkpoints

The potential functional relationships between ICs were analyzed and visualized as a network using STRING (Search Tool for the Retrieval of Interacting Genes/Proteins; string-db.org/), a robust database and computational resource that offers comprehensive information on protein‒protein interactions from diverse sources, including experimental data, computational predictions, and literature^32^.

### Uniform Manifold Approximation and Projection (UMAP)

The UMAP analysis was conducted using the ‘umap’ function in R and standardized data. The default configuration was set with a n_neighbors value of 15, n_components set to 2, and min_dist set at 0.1. UMAP was chosen for its ability to capture the nuances of high-dimensional data and effectively project them onto a two-dimensional space, highlighting potential clusters or patterns.

### Principal Component Analysis (PCA)

PCA is sensitive to the scale of variables and performs best with normally distributed data. Variables vastly different in the order of magnitude can lead to biased principal components that reflect the larger scale variables more. However, transcriptomic data often exhibit a wide range of abundance values and are not normally distributed. To address these issues, the data were first log2-normalized. This transformation compressed the scale of the data and approximated a normal distribution. Subsequently, the normalized data were standardized (centered and scaled) within the ‘prcomp’ or ‘PCAtools::pca’ R functions. This step involved subtracting the mean and dividing by the standard deviation of each variable, which ensured that all variables contributed equally to the analysis. Next, PCA was performed with the aforementioned functions.

### Comparison of healthy vs. cancerous tissue

Healthy reference samples originated from areas adjacent to primary tumors^21^.

### Computational environment

Below, detailed information on the computational environment used for this study is presented. The raw data preprocessing and read quantification were performed on a Prometheus cluster, and the detailed data analysis was conducted in R 4.2.3 (2023-03-15 ucrt) using RStudio 2022.07.0 (Build 548) on the platform x86_64-w64-mingw32/x64 (64-bit) running under Windows 10 x64 (build 22621). Detailed information about the versions of all directly used R packages and their recursively defined dependencies is provided in **Table S7**.

## RESULTS

### Immune Checkpoints Are Expressed Differently Across Canine Cancer Types

Diving into the largely uncharted territory of canine cancer immunology, we set out to explore whether spontaneous canine cancers express immune checkpoints (ICs) relevant to human cancers. To probe this, we harnessed publicly accessible raw RNA sequencing data^33–45^. This yielded a robust dataset from 418 canine patients spanning 14 distinct cancer types (**Tab. S5**). Additionally, we compiled a comprehensive list of both established and emerging ICs (**Tab. S6**). We quantified gene-level expression and visualized the normalized median abundances of those IC genes in **Fig. 1A**, along with their impact on the immune response and best known interactions. Importantly, all ICs were expressed in at least some canine cancers. While SIRPA-CD47 and TIM-3-GAL-9 inhibitory receptor‒ligand pairs were universally highly expressed, the abundance of others, like B7-H4 (inhibitory), NECTIN4 (inhibitory) and GITR (complex role), varied greatly. Interestingly, in the case of the commonly drugged PD-1/PD-L1 inhibitory pair, oral squamous cell carcinoma (OSCC) and meningioma exhibited low PD-1 receptor expression, while OSCC displayed elevated PD-L1 ligand levels alongside undetectable PD-1. The distribution of normalized abundances for each IC across canine cancer types can be found in **Fig. S4**. The standard deviation and median absolute deviation/median (MADm) plot to match Fig. 1A is available as **Fig. S2**.

**Figure 1:**
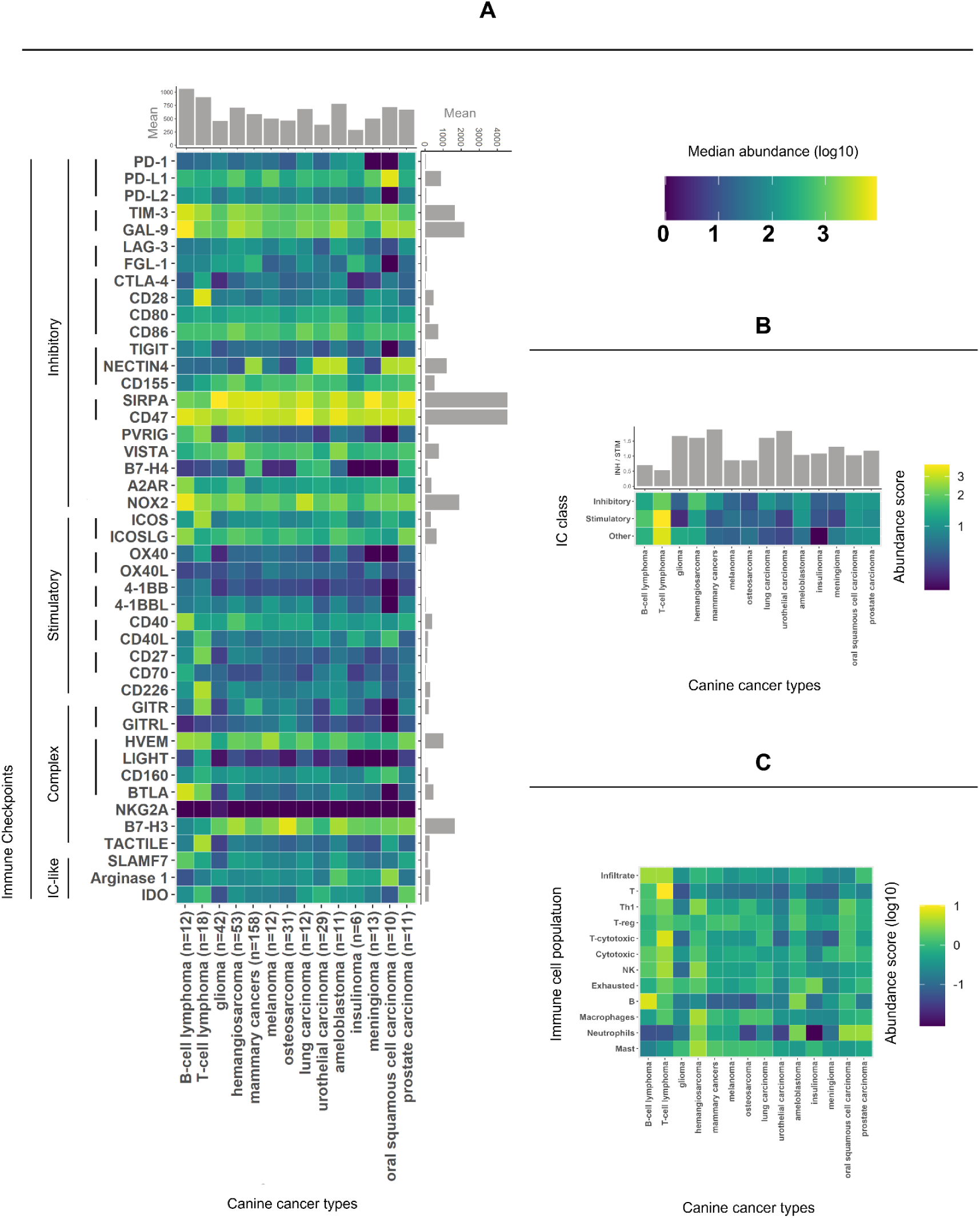
The immune environment across canine cancer types. (A) Normalized median abundances of immune checkpoints (ICs). Notably, ICs such as TIM-3 and GAL-9 are universally expressed, while others, such as GITR, B7-H4, and NECTIN4, are variable. The gray accessory plots summarize the gene expression per cancer and per IC, with lymphomas showing the strongest total IC expression and SIRPA/CD47 exhibiting the highest total expression. (B) This heatmap displays the abundance scores for the IC categories. The bar plot above illustrates the ratio between inhibitory and stimulatory scores, with a ratio of 1 indicating an equal balance. Notably, MC, UC, LC, GLM, and HSA (see abbreviations) had high ratios, suggesting a potentially more inhibitory environment. (C) A heatmap of standardized scores for various immune cell populations showing the diversity in immune infiltration across different cancer types.

### The Extent of Immune Inhibition Varies Across Canine Cancer Types

How does the IC landscape of each cancer influence resident immune cells? Single ICs such as PD-1 do not reflect the general level of immune inhibition within the tumor microenvironment^46^. This complexity is underscored by the compensatory role of other ICs, such as TIM-3/GAL-9, which can drive resistance to PD-1 blockade immunotherapy^47^. In an attempt to assess the level of immune inhibition in the tumor environment, we categorized the ICs under study into inhibitory, stimulatory, or complex groups and summarized the abundance of each category into inhibitory and stimulatory IC scores (see Methods section - calculating the immune inhibition score). We then computed the ratio of the inhibitory score to the stimulatory score (**Fig. 1B**). MC, UC, LC, GLM, and HSA exhibited a notably high ratio (a more inhibitory environment), while BCL, TCL, MEL, and OS exhibited low values.

### Diverse Immune Infiltrates Characterize Canine Cancers

To contextualize the abundance of ICs within each cancer’s environment, we evaluated the expression of immune cell markers. Advanced methods for deconvoluting immune infiltrates from sequencing data, such as CiberSort, were not yet available for canine model; hence, we quantified the expression of marker genes characteristic of each cell population and summarized them as population estimate scores (**Fig. 1C**; see methods: immune infiltrate estimation). We found a particularly high estimated immune infiltration in prostate carcinoma and lymphomas (artifact expected due to the immune nature of lymphoma cells). The estimated scores for T- and cytotoxic T-cells were high in T-cell lymphoma, while the scores for B-cells were high in B-cell lymphoma, which was expected due to the nature of these cancers. Additionally, the B-cells score appeared to be high in ameloblastoma. Macrophage and mast cell scores were high in hemangiosarcoma. Neutrophil score appeared high in hemangiosarcoma, ameloblastoma, prostate carcinoma and oral squamous cell carcinoma. Additionally, we observed a relatively high score for exhausted immune cells in insulinoma. Glioma appeared low in most immune cell types save for T-reg and exhausted lymphocytes. Intriguingly, glioma had a particularly low score for NK cells, while being the only canine cancer with a considerable abundance of NKG2A, the NK cell receptor (**Fig. 1 A & C, Fig. S4**). Other low scores included neutrophils in osteosarcoma, urothelial carcinoma and especially insulinoma. Our findings reveal a broad variation in the estimated composition of immune infiltrates across canine tumors.

### Cancer IC Signatures Cluster by Species

After discovering variations in the immune environment of canine tumors, we sought to determine whether these canine cancers mirror their human counterparts in terms of immunotherapy targets, specifically immune checkpoints. Accordingly, we compared cancer IC signatures within and between human and canine species. We combined our data on IC abundance in canine cancer types (n=14, 418 patients) with human cancer data (n=27, 1200 patients) supplied by the PanCancer Analysis of Whole Genomes (PCAWG) and sourced from EBI^21^. Subsequently, we quantified the median abundance of each IC in each cancer type and created a vector of these medians for every cancer type (see methods). We assessed the similarity of these vectors using a Pearson correlation-based distance matrix. The dendrogram resulting from the hierarchical clustering of the matrix showed that cancer types predominantly clustered within their respective species, as shown in **Fig. 2**. In other words, when cancers were grouped based on IC expression similarity, canine cancers primarily clustered with other canine types, while human cancers largely grouped together as well.

**Figure 2:**
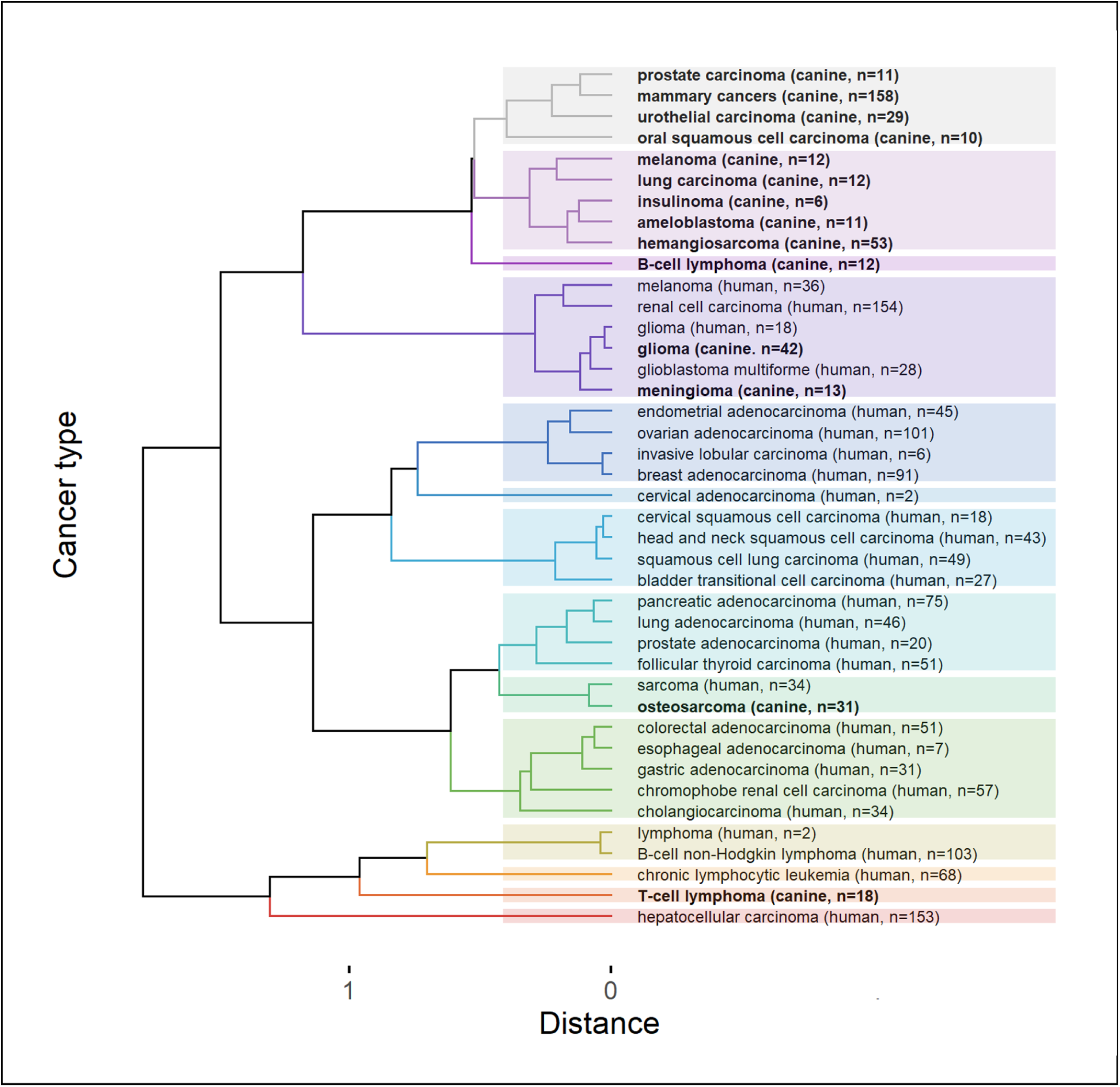
Dendrogram representing the hierarchical clustering of a distance matrix that compares canine and human cancers with respect to their immune checkpoint (IC) expression levels. Cancers grouped predominantly within their respective species (**bold** font - canine, normal font - human). However, notable interspecies similarities emerged, particularly between canine and human gliomas and between canine osteosarcoma and human sarcoma. The figure also illustrates that human cancers tend to cluster according to their histological subtype and primary cancer site. Colors distinguish 14 different clusters. For detailed distance values, refer to **Table S4**.

### Transcending Species Lines: Brain Cancers and Sarcomas

Remarkably, there were two interspecies groups that did not follow the trend of clustering within species. Firstly, canine glioma and meningioma, along with human glioma and glioblastoma, displayed striking relatedness. Our analysis of the distance table (**Tab. S4**) revealed that the interspecies glioma distance (0.025) was remarkably smaller than any distance found within the same species for humans or dogs, where the lowest recorded distances were 0.032 and 0.083, respectively. Secondly, canine osteosarcoma and human sarcoma demonstrated exceptional correlation, even though sarcoma represents a broad category (0.086). Delving further into **Tab. S4**, we observed that many human-canine pairs displayed distances that were lower than the median intraspecies distances (0.344 for dogs and 0.541 for humans, shown in **Tab. S4**, violet and green), thus illustrating broad similarity.

### In Human Cancers, IC Signatures Reflect the Histology and Primary Site

For a more granular view of type-specific patterns, we inspected human cancers, as these were categorized into more precise diagnostic subtypes. Human cancers appeared to cluster by histological type and subtype (e.g., distinct clustering of squamous cell carcinomas vs adenocarcinomas). Clustering also appeared to follow physiologically related primary sites, evident in the grouping of endometrial, ovarian, invasive lobular and breast adenocarcinomas vs. gastric, esophageal, and colorectal adenocarcinomas (**Fig. 3**).

**Figure 3:**
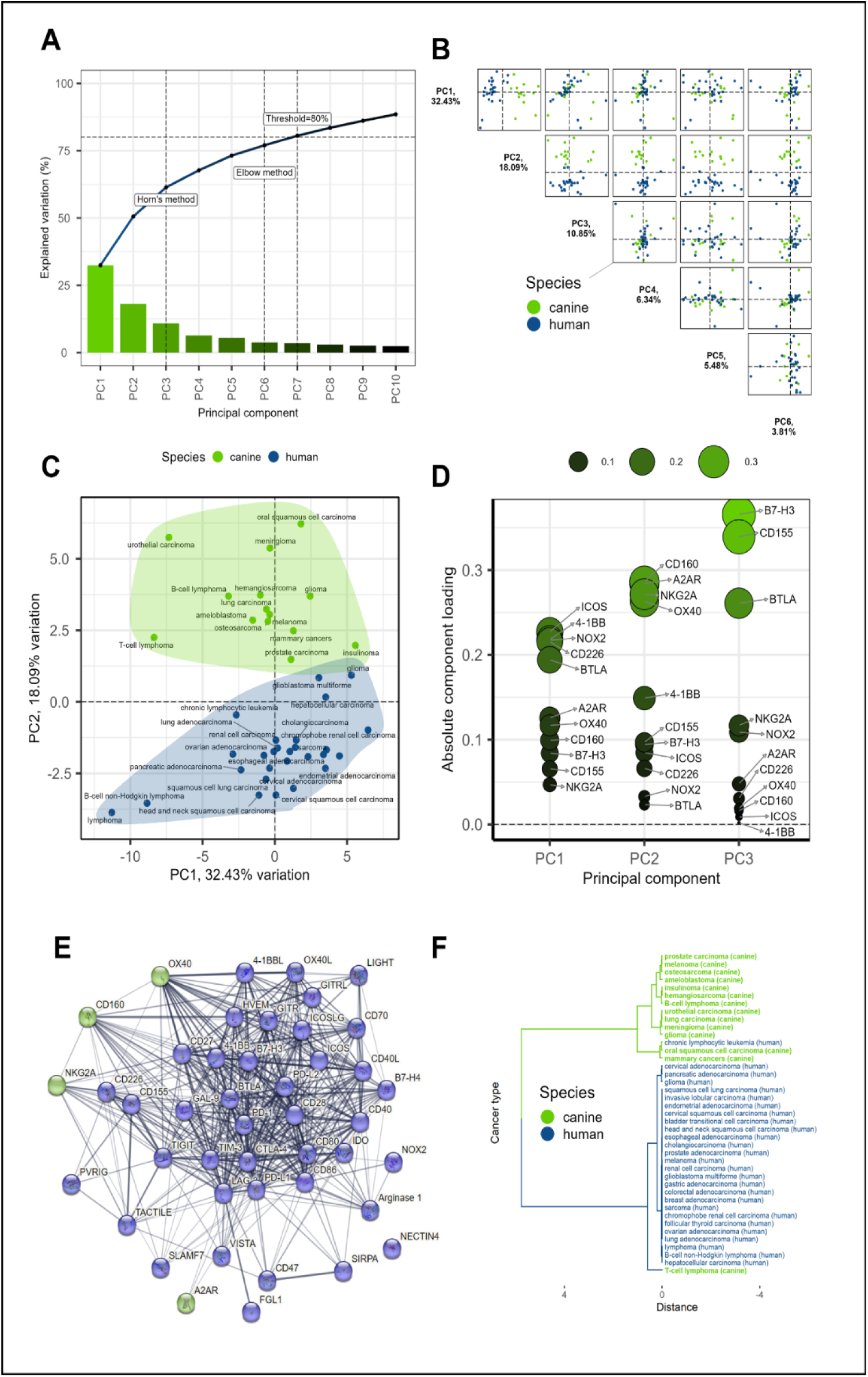
Principal component analysis (PCA) of immune checkpoint (IC) signatures in human and canine cancers. (A) Optimal dimensionality determination for PCA using the elbow method. (B) Visualization of the top five principal components (PCs). (C) Clear species separation along the second principal component (PC2). (D) Component loadings for PC1 to PC3, highlighting four influential ICs (CD160, A2AR, NKG2A, and OX40) contributing to PC2. (E) StringDB-based analysis showing that immunoregulatory function was the only feature shared among the four influential ICs. (F) Dendrogram resulting from clustering of signatures consisting solely of the 4 identified genes, demonstrating nearly complete separation of cancers by species.

To determine whether these clustering tendencies were driven by the abundances of any specific ICs, we investigated median-based IC differential expression between the clusters of interest and the remaining cancer types or between particular clusters. The statistically significant results defined by p < 0.05 were sorted by decreasing fold change and are presented in **Table 1** below. Additionally, the outcomes of the mean-based analysis are shown in **Table S1**. The significant ICs found in both analyses are marked in bold.

**Table 1:** Median-based differential IC expression between human cancer type clusters. Genes that were found to be significant in both the median- and mean-based analyses are marked in bold to indicate increased confidence. Fc - fold change, log2fc - Log of fc, pval - p value, HCC - hepatocellular carcinoma, MEL - melanoma, RCC - renal cell carcinoma, SCC - squamous cell carcinoma, TCC - transitional cell carcinoma.

| Comparison | ID | median tpm | reference tpm | fc | log2_fc | zscore | pval | regulation |
| --- | --- | --- | --- | --- | --- | --- | --- | --- |
| Sarcoma vs other | **NECTIN4** | 0.40 | 22.50 | 0.02 | -5.81 | -4.01 | 2.9894E-05 | down |
|  | **B7-H4** | 0.10 | 1.50 | 0.07 | -3.91 | -2.63 | 4.2311E-03 | down |
|  | CD70 | 6.00 | 1.00 | 6.00 | 2.58 | 2.07 | 1.9363E-02 | up |
| Melanoma vs other human | **NECTIN4** | 0.50 | 22.50 | 0.02 | -5.49 | -4.30 | 8.5662E-06 | down |
|  | **B7-H4** | 0.10 | 1.50 | 0.07 | -3.91 | -3.02 | 1.2592E-03 | down |
| HCC vs other | **FGL-1** | 530.00 | 0.10 | 5300.00 | 12.37 | 4.23 | 1.1564E-05 | up |
|  | **Arginase 1** | 335.00 | 0.20 | 1675.00 | 10.71 | 3.68 | 1.1883E-04 | up |
|  | **NECTIN4** | 0.20 | 22.50 | 0.01 | -6.81 | -2.20 | 1.3918E-02 | down |
| Gynecological vs other | **B7-H4** | 86.50 | 1.50 | 57.67 | 5.85 | 5.30 | 5.9158E-08 | up |
| Gastric cluster vs other | **B7-H4** | 0.40 | 1.50 | 0.27 | -1.91 | -3.14 | 8.3576E-04 | down |
|  | **NECTIN4** | 10.00 | 22.50 | 0.44 | -1.17 | -1.88 | 3.0173E-02 | down |
|  | **GAL-9** | 68.00 | 31.50 | 2.16 | 1.11 | 2.04 | 2.0919E-02 | up |
|  | **CD70** | 2.00 | 1.00 | 2.00 | 1.00 | 1.85 | 3.2445E-02 | up |
| Lymphomas vs other | **BTLA** | 29.00 | 0.40 | 72.50 | 6.18 | 1.66 | 4.8874E-02 | up |
|  | **NECTIN4** | 0.50 | 22.50 | 0.02 | -5.49 | -3.24 | 5.8839E-04 | down |
|  | **CD155** | 4.50 | 36.50 | 0.12 | -3.02 | -2.21 | 1.3671E-02 | down |
|  | **B7-H3** | 7.00 | 55.00 | 0.13 | -2.97 | -2.19 | 1.4359E-02 | down |
| Brain cancers vs other | **NECTIN4** | 0.15 | 22.50 | 0.01 | -7.23 | -3.17 | 7.5006E-04 | down |
|  | **IDO** | 0.20 | 7.50 | 0.03 | -5.23 | -2.12 | 1.7039E-02 | down |
| Brain cancers cluster with MEL and RCC vs other | **NECTIN4** | 0.20 | 22.50 | 0.01 | -6.81 | -5.08 | 1.8943E-07 | down |
|  | **SIRPA** | 153.50 | 48.50 | 3.16 | 1.66 | 1.68 | 4.6889E-02 | up |
| SCC and TCC vs other | **GITR** | 27.00 | 4.50 | 6.00 | 2.58 | 2.78 | 2.6858E-03 | up |
|  | **NECTIN4** | 105.50 | 22.50 | 4.69 | 2.23 | 2.33 | 9.9524E-03 | up |
|  | **B7-H4** | 6.00 | 1.50 | 4.00 | 2.00 | 2.03 | 2.0954E-02 | up |
| Carcinomas vs other | **B7-H4** | 7.00 | 1.50 | 4.67 | 2.22 | 5.09 | 1.7518E-07 | up |
|  | **NECTIN4** | 43.50 | 22.50 | 1.93 | 0.95 | 2.05 | 2.0195E-02 | up |
| Adenocarcinomas vs other | **B7-H4** | 9.00 | 1.50 | 6.00 | 2.58 | 4.61 | 2.0487E-06 | up |
|  | **ICOS** | 2.00 | 0.90 | 2.22 | 1.15 | 1.77 | 3.8458E-02 | up |
| Gynecological adenocarcinomas vs other adenocarcinomas | **IDO** | 30.50 | 9.00 | 3.39 | 1.76 | 3.89 | 4.9942E-05 | up |
| Gastric adenocarcinomas vs other adenocarcinomas | **B7-H4** | 0.40 | 66.00 | 0.01 | -7.37 | -5.73 | 4.9198E-09 | down |

Uncharacteristically, the levels of Nectin4 and/or B7-H4 were significant in almost all clusters. Carcinomas were characterized by upregulated B7-H4 and NECTIN4 as compared to non-epithelial cancers. SCC and TCC carcinomas, which clustered together, additionally displayed elevated GITR. On the other hand, another type of carcinoma, adenocarcinomas, distinguished itself by slightly elevated ICOS. Within adenocarcinomas, we assessed two separate clusters: gynecological and gastric ones. Compared to all adenocarcinomas, the gynecological ones featured upregulated IDO, while gastric adenocarcinomas had severely downregulated B7-H4. Brain cancers had down-regulated NECTIN4 - consistent with their non-epithelial character - and down-regulated IDO. When the whole cluster was considered - glioma and glioblastoma with their close HC neighbors melanoma and renal cell carcinoma, elevated SIRPA became a statistically significant feature. Down-regulation of Nectin4 and B7-H4 was the strongest characteristic of human sarcoma vs all other human cancers. However, the same was true for melanoma. This highlights the limitation of differential-expression based methods. The expression analysis of individual genes does not identify features unique to clusters identified with more comprehensive methods. Arginase 1 and FGL-1 levels were extremely elevated in hepatocellular carcinoma, as they are in healthy liver tissues^48^.

### A2AR and Six Other ICs Are Possible Drivers of the Species Divide

What were the roots of the differences between human and canine IC signatures? Are these distinctions due to complex, species-specific characteristics, or are a handful of IC genes the main influencers? To address these questions, we first performed a differential IC expression analysis - as above - comparing the human and canine cancer type clusters (**Tab. 2** - below). Five ICs appeared to distinguish canine from human cancer types.

**Table 2:** Median-based differential IC expression comparing human vs canine cancer types. Genes that were found to be significant in both the median- and mean-based analyses are marked in bold to indicate increased confidence. Fc - fold change, log2fc - Log of fc, pval - p value.

| Comparison | ID | median tpm | reference tpm | fc | log2_fc | zscore | pval | regulation |
| --- | --- | --- | --- | --- | --- | --- | --- | --- |
| Canine vs Human | **FGL-1** | 4.78 | 0.10 | 47.75 | 5.58 | 2.61 | 4.5128E-03 | up |
|  | **Arginase 1** | 5.21 | 0.20 | 26.04 | 4.70 | 2.18 | 1.4554E-02 | up |
|  | **NECTIN4** | 0.77 | 18.00 | 0.04 | -4.55 | -2.36 | 9.2459E-03 | down |
|  | **A2AR** | 4.27 | 0.20 | 21.33 | 4.41 | 2.04 | 2.0641E-02 | up |
|  | **OX40L** | 0.32 | 3.00 | 0.11 | -3.24 | -1.71 | 4.3182E-02 | down |

However, only the A2AR was indicated in both the median- and mean-based analyses (**Tab. 2 & S1**). Subsequently, we decided to answer the question of species division by performing a principal component analysis (PCA) on the IC signatures (refer to **Fig. 3**).

Principal Component Analysis (PCA) is a method that reduces the complexity of high-dimensional data while retaining key patterns. In the original dataset, each variable or sample can be considered a dimension. PCA generates as many ‘principal components’ (PCs) as there are dimensions - 44 in our case because there are 44 genes (variables) but only 41 cancers (samples). These PCs are essentially new ‘dimensions’ that are combinations of the original ones. However, PCs are ranked (PC1-PC44) based on their contribution to the overall variability. The first few PCs usually encapsulate a significant portion of this variability; thus, by focusing on these PCs, we simplify further analyses and enable meaningful data visualization.

We determined the optimal number of PCs to retain using a scree plot, which visualizes the proportion of the total data variance that is captured by each PC (**Fig. 3A**). We visualized the top 10 PCs, marked the threshold capturing at least 80% of the data variability, and analyzed the optimal number of PCs to consider with Horn’s and Elbow methods. The elbow method identifies an ‘elbow’ point at which the addition of more PCs provides diminishing returns in terms of explained variability. Based on the more conservative elbow estimate, we chose to retain the 6 top PCs.

Following the selection of the top six PCs, we proceeded to investigate their interplay (**Fig. 3B**). Studying combinations of PCs, rather than each individually, provides a more comprehensive understanding of the dataset, as it allows us to explore the simultaneous effect of multiple components on variability. Strikingly, PC2 alone clearly separated the human and canine groups (**Fig. 3C**).

For deeper insight, we scrutinized the ‘component loadings’ for PC1 to PC3 (**Fig. 3D**). Component loadings represent the weight by which each initial variable (in this case, each gene) contributes to the principal component. By investigating these loadings, we can identify the key players driving the differentiation captured by each PC. We pinpointed the four most influential ICs contributing to PC2, while noting that these ICs were not the primary contributors to the variability in PC1 and PC3 (**Fig. 3D**).

These ICs - CD160, A2AR, NKG2A, and OX40 - were further examined for potential enrichment in various categories using StringDB, but they appeared to share only their immunoregulatory function. We visualized the StringDB-based relationships between all the analyzed IC genes (**Fig. 3E**), which clarified the weak links between the four genes of interest. Subsequently, we repeated hierarchical clustering as before but with signatures consisting solely of the four identified genes or only four random genes for comparison (VISTA, ICOSLG, 4-1BBL, TACTILE). Remarkably, clustering the signatures consisting solely of the 4 identified genes resulted in a nearly complete separation of cancers by species (**Fig. 3F**), while clustering by 4 random genes allowed the cancer types of both species to intermingle freely (**Fig. S7**). Broadly speaking, in canine versus human cancers, the abundances of CD160 and A2AR were greater, while those of NKG2A and OX40 were lower (**Tab. S2**). Importantly, these trends should not be assumed to apply universally to all individual cancer cases or cancer types, as exemplified by canine glioma NKG2A expression, isolated amongst canine malignancies (**Fig. S4**).

Summarizing the results of the two experimental approaches, our confidence is predominantly anchored in the role of A2AR as a determinant of the interspecies divide. Given the very low median values for NKG2A in both species (see **Tab. S2**) and considering that the majority of canine expression is attributable to glioma alone, we infer that the influence of NKG2A is likely statistical rather than biological. The quartet of ICs identified through differential expression analysis did not converge with the two other ICs isolated by PCA, a finding consistent with the disparate nature of these methodologies. This discrepancy, coupled with the constraints of TPM normalization used in the input data, limits our confidence in these six other ICs. In conclusion, the species divide appears to be primarily driven by A2AR, possibly in concert with one or more of the following immune checkpoints: FGL-1, Arginase 1, NECTIN4, OX40L, CD160, OX40. From a comparative oncology perspective, immune checkpoints with notable abundance variability within or between species may not be optimal treatment targets.

### Key IC Genes Driving Non-Species Differences in Cancer Types

PCA also revealed that the IC genes ICOS, 4-1BB, NOX2, CD226 and BTLA had the highest loadings on PC-1 and not on PC-2 (**Fig. 3D**). Between PC-1 and PC-2, 4-1BB, an inducible costimulatory IC receptor of T-cells, was the only one unique to PC-1. Given that PC-1 captures the most variability in the dataset, the abundances of these five ICs, especially 4- 1BB, may contribute the most to the non-species-related differences among the analyzed cancer types. 4-1BB is considered a promising cancer immunotherapy target^49^.

### Case-by-Case: Individual Tumors Unveil IC Signature Heterogeneity

At the outset, we used gene expression medians to discern patterns across cancers. To determine the internal heterogeneity of IC signatures within each cancer type, we decided to turn our attention to individual cases. We sought to determine whether cases of the same cancer type share associated IC signatures or if the signatures are unique to each case, irrespective of the diagnosis. We were also interested in outliers that would defy their classification by aligning with cases of other cancer types. We processed individual IC abundance signatures with UMAP for its ability to capture the complex and potentially non-linear relationships among the expression levels of analyzed genes in each patient’s profile. This is important because gene expression levels often interact in intricate ways that linear methods can overlook. The use of UMAP allowed us to maintain both broader patterns and finer details in our data (**Fig. 4**). This method is especially effective for revealing significant insights into individual cases and outliers where nuanced patterns can have substantial implications. Our findings echo previous observations at the cancer type level; individual canine and human cancer cases formed distinguishable clusters (**Fig. 4A**). Gliomas in both species were closely situated, demonstrating highly uniform signatures (**Fig. 4B**). In fact, all brain cancer cases presented a high degree of proximity (**Fig. 4C**). Sarcomas in both species also exhibited clustering, albeit less tight (**Fig. 4D**). Hemangiosarcoma, a predominantly canine cancer, did not cluster as closely with human sarcomas, as did canine osteosarcoma. Certain cancer types, such as human cholangiocarcinoma, presented a high degree of dispersion in their individual case signatures, thereby colocalizing with other cancers (**Fig. 4E**). This precludes meaningful interpretation of averaged trends for such cancer types. In line with earlier hierarchical clustering, certain human canine cancer sets, such as prostate carcinomas, did not exhibit similar signatures (**Fig. 4F**).

**Figure 4:**
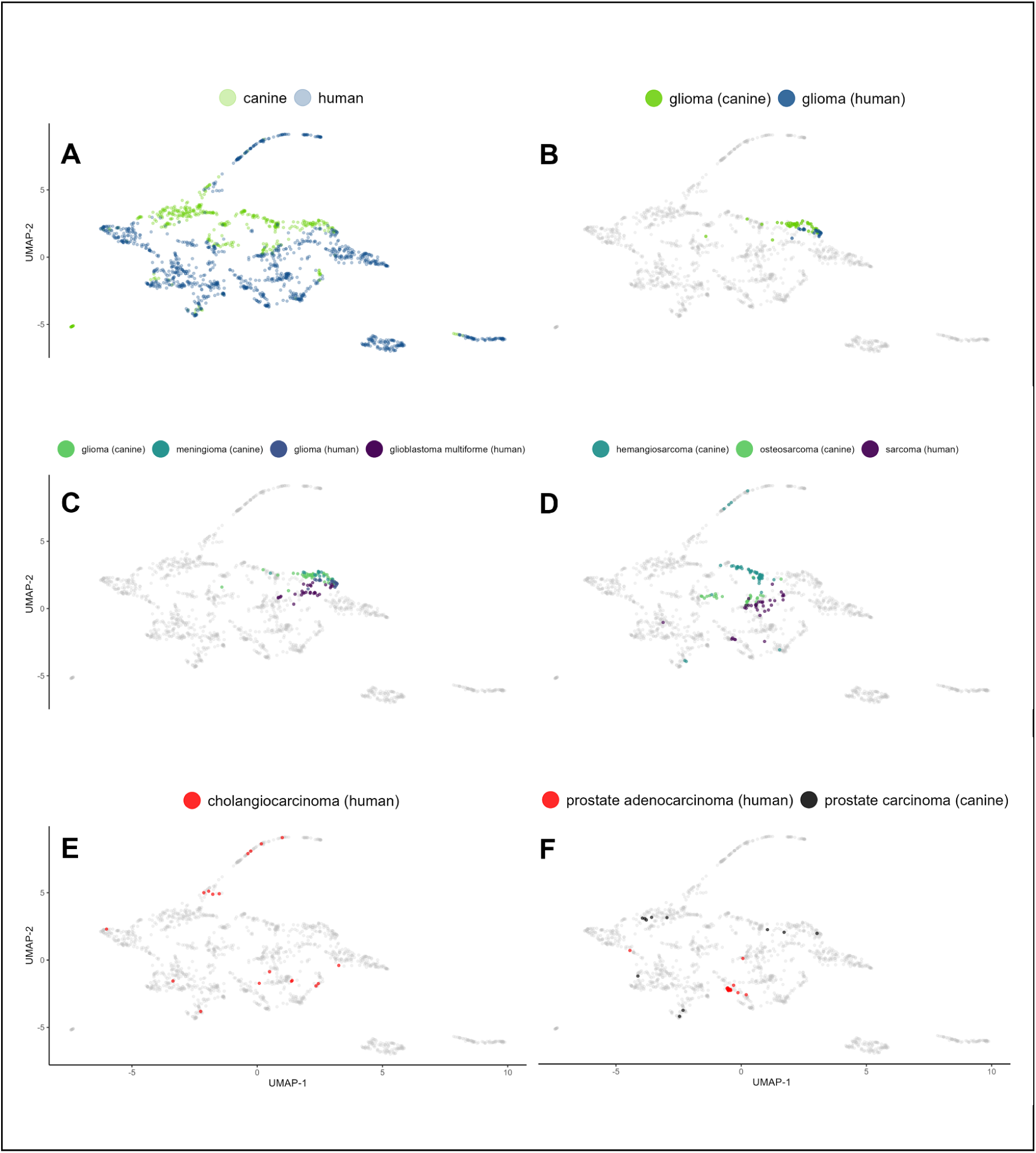
Individual IC abundance signatures via UMAP representation underscore cancer type-specific patterns and intrinsic heterogeneity. (A) Global overview of the signatures distinguishes two primary clusters: canine and human, highlighting species-dependent variation. (B) Glioma cases from both species are closely situated, signifying a shared IC signature landscape. (C) Inspection of brain cancer cases from both species reveals not only mutual proximity but also marked uniformity within each cancer type. (D) Examination of sarcoma cases unveils a proximity between human sarcoma and canine osteosarcoma, while canine hemangiosarcoma exhibits a more distant relationship. (E) Human cholangiocarcinoma displays notable case dispersion, indicative of a high degree of heterogeneity in the IC signature. (F) A cross-species comparison of prostate cancer evidences a lack of coclustering, suggesting distinct IC signatures for these species-specific prostate cancers.

### Confirming Patterns and Unveiling Surprises: Revisiting Patient Data with PCA

To complement our UMAP analysis, we applied PCA to the same dataset, providing a linear perspective to validate and expand the non-linear UMAP insights. **Figure 5** displays PCA plots of the log-normalized IC signature data of individual patients. The consistency of the primary trends between UMAP and PCA strengthens our findings. **Figure 5A** provides a global overview of the PCA analysis, distinguishing two primary clusters representing canine and human cases. The plot reinforces the species-dependent variation observed in the UMAP analysis and in the earlier cancer type-focused PCA. In line with the UMAP results, the PCA confirmed that brain cancers, particularly gliomas, of both species exhibited close proximity (**Fig. 5C**), indicating a shared immune checkpoint signature landscape. Sarcomas, characterized by a less homogeneous distribution of patient signatures, demonstrated more separation along the species-distinguishing PC-2 axis but still exhibited a reasonably similar distribution along the PC-1 axis, which captured the most variability in the dataset (**Fig. 5D**). Prostate carcinomas, on the other hand, did not appear well aligned along the PC-1 axis, confirming their distinct IC signatures and serving as an example of a human-dog cancer pair with low potential for comparative immunotherapy (**Fig. 5F**). Similar to UMAP, PCA confirmed that human cholangiocarcinoma displayed a wide distribution of data points along the PC-1 axis, reflecting a high degree of intratumoral heterogeneity in the IC signature (**Fig. 5E**). The PCA also revealed a novel finding: a cluster of human patients with signatures uniquely positioned within the plot area belonging to the canine cancer patients (**Fig. 5B**). Notably, all these patients had human chronic lymphocytic leukemia (CCL), a B-cell-based blood cancer. Although no equivalent leukemia was present in our canine dataset for direct comparison, this observation suggests unique resemblance between the IC landscapes of human CCL and canine malignancies (**Fig. 5B**). In the earlier UMAP analysis, CCLs formed a very distinct cluster neighboring only some canine B-cell lymphoma cases (**Fig. S6**). CCL was the only leukemia included in our study, and it is important to acknowledge that the differential sample preparation methods employed for leukemia versus solid tumors could have influenced the observed results. Nevertheless, this unexpected finding highlights the value of multimethod analysis and warrants future exploration. In summary, the PCA validated and expanded upon the UMAP findings.

**Figure 5:**
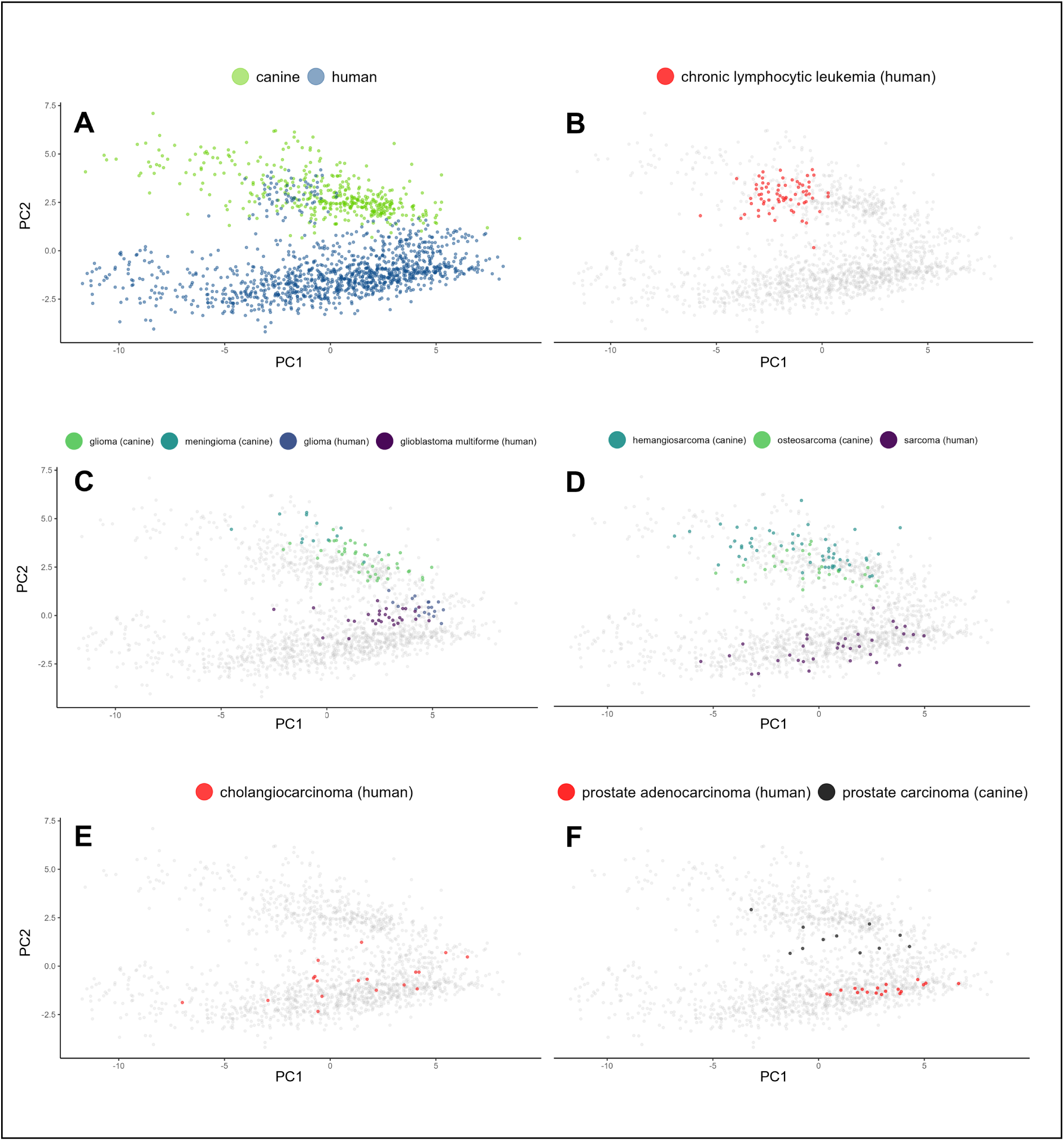
PCA representation of individual IC signatures validates and augments the insights from UMAP analysis, revealing unexpected similarities between human chronic lymphocytic leukemia (CCL) and canine cancers. (A) Consistent with the UMAP findings, the global overview reveals two primary clusters: canine and human. (B) Notably, the PCA revealed a surprising finding missed by UMAP: patient signatures of human CCL, a B-cell blood cancer, exhibit a striking similarity to canine cancers along the PC2 axis, which accounts for between-species differences. This finding suggests a resemblance between human CCL and canine malignancies in the context of the identified key immune checkpoint genes. (C) Further examination of brain cancer cases reaffirms the mutual proximity observed in UMAP, which is particularly evident in gliomas, and the overall uniformity of signatures within each cancer type. (D) Human sarcoma and canine osteosarcoma, while displaying variability along PC-1, demonstrate alignment with each other, further supporting the concordance between species. (E) Similar to UMAP, human cholangiocarcinoma exhibits notable case dispersion, reflecting the high degree of heterogeneity in IC signatures. (F) Consistent with previous observations, PCA confirmed that human and dog prostate carcinomas display distinct IC signatures, as they do not align along PC-1.

### Distinguishing Cancer and Healthy Tissue Through IC Expression Patterns

Our primary goal was to compare IC signatures between various cancers, both within and across species. Thus, we concentrated on the baseline IC expression in cancer rather than the differential expression between cancerous and healthy tissues. Regardless of the IC status in healthy tissues, the target abundance in cancer alone offers valuable insights for immunotherapy. However, five human cancers had at least nine healthy references available, so we took the opportunity to evaluate whether cancer IC signatures mirror the characteristics of their healthy tissues of origin or whether they exhibit an independent character. To this end, we visualized the diseased samples compared to healthy adjacent tissues, as assessed by UMAP and PCA. This time, the analyses were extended to include all 150 normal samples present in the PCAWG dataset (**Fig. S3**).

In UMAP, the small sample size limited the interpretations for cholangiocarcinoma and hepatocellular carcinoma, but renal cell carcinoma and chromophobe renal cell carcinoma presented a very distinct shift between the normal and cancerous samples (**Fig. 5A-E**). Lung adenocarcinoma also exhibited a notable contrast in the immune checkpoint profiles of tightly clustered normal samples and the more diverse set of cancer samples. It’s important to acknowledge that the ‘normal’ reference samples from the PCAWG RNA-seq study were obtained from areas adjacent to primary tumors^21^. The potential sharing of features between tumor and nearby stroma could downplay the observed differences.

The PCA mirrored these findings (**Fig. 7A-E**), particularly with the renal carcinomas demonstrating an observable shift in the distribution of healthy and cancer samples along the PC-1 axis. CD86, TIGIT, ICOS, and BTLA emerged as genes with the greatest contributions to the PC-1 component loadings (**Fig. 7F**). As such, these IC genes merit further exploration for their possible differential expression in cancer and their potential as immunotherapy targets in renal cell carcinomas, which are currently treated with PD-1 and CTLA-4-blocking therapeutics^50^.

**Figure 6:**
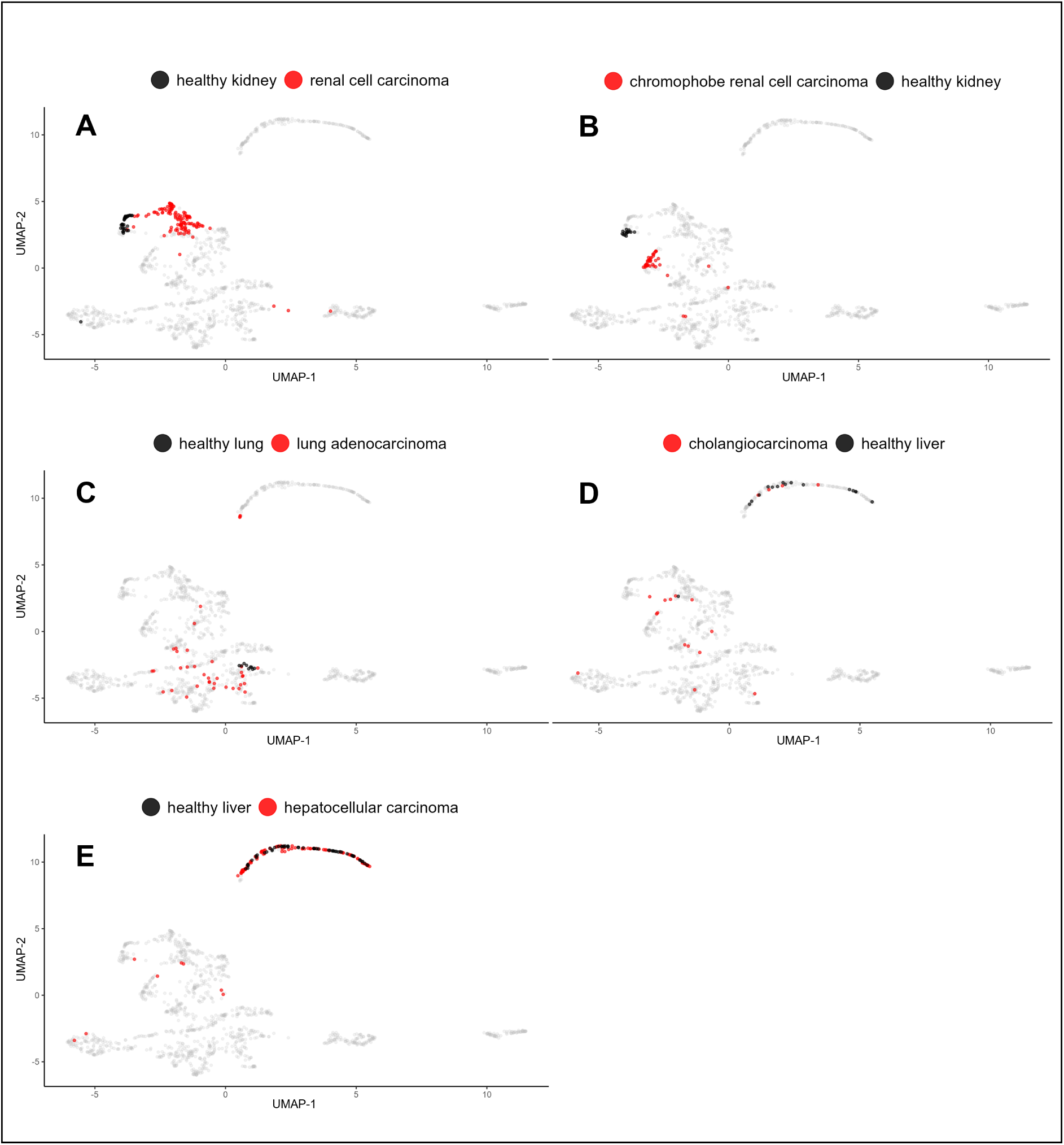
Immune Checkpoint Expression Patterns Differ Between Cancers and Their Corresponding Normal Samples. UMAP visualizations of IC gene expression patterns in individual samples of five different cancer types along with corresponding healthy tissues. (A) Renal cell carcinoma, (B) chromophobe renal cell carcinoma, (C) lung adenocarcinoma, (D) cholangiocarcinoma, and (E) hepatocellular carcinoma. Distributions provide insights into differential IC landscapes between cancerous and normal tissues, with a notable shift in the case of renal carcinomas and lung adenocarcinoma.

**Figure 7:**
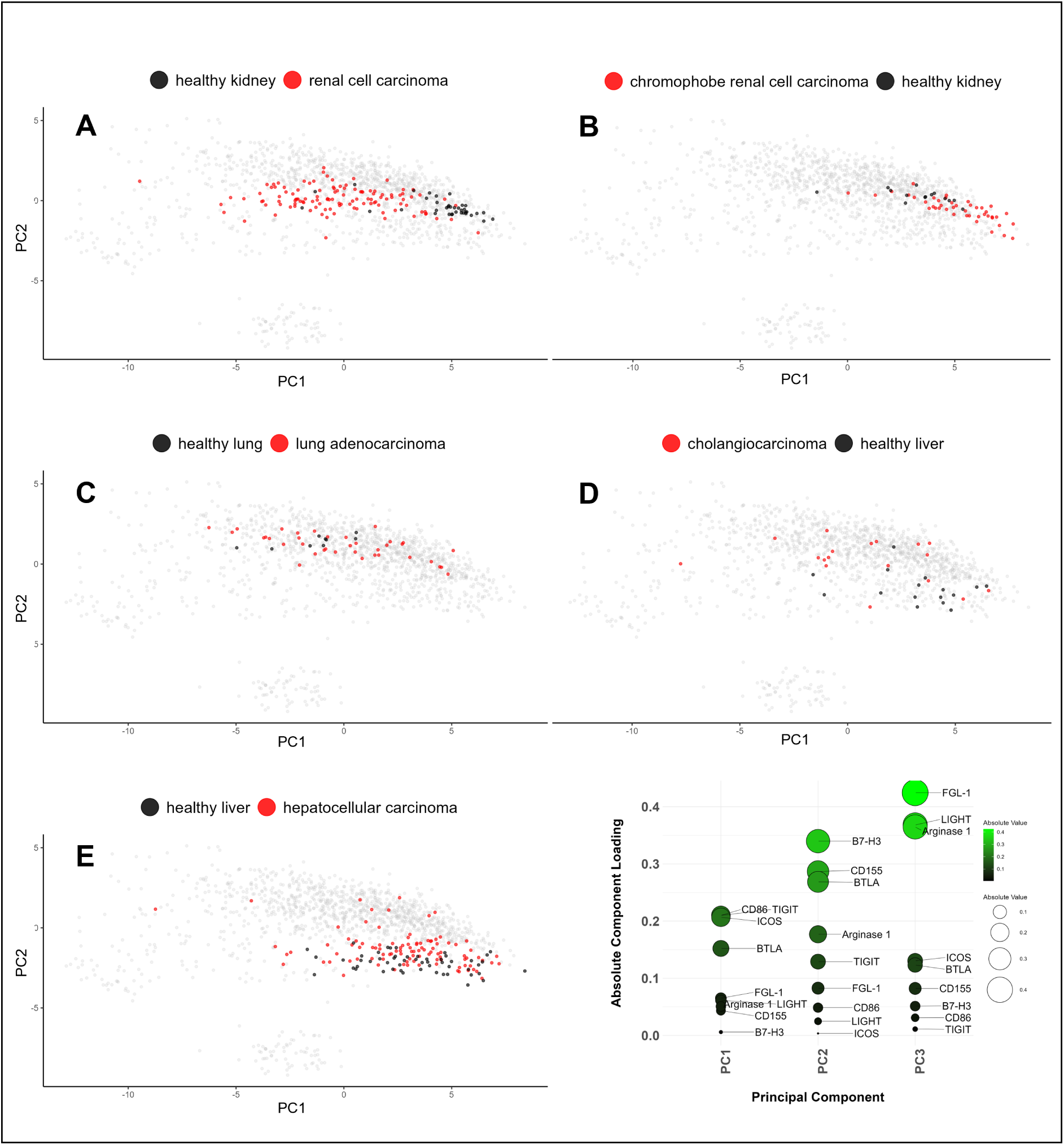
Differential Immune Checkpoint Expression Patterns Between Cancers and Normal Samples Unveiled Through PCA and Component Loadings Analysis. PCA biplots (PC-1 vs PC-2) present the IC expression distributions for the same cancer types and corresponding healthy tissues as in Figure 6. (A) Renal cell carcinoma, (B) chromophobe renal cell carcinoma, (C) lung adenocarcinoma, (D) cholangiocarcinoma, and (E) hepatocellular carcinoma. (F) A plot of component loadings for PC-1 to PC-3, emphasizing the genes CD86, TIGIT, ICOS, and BTLA, which show the most substantial contributions to the PC-1 component. These genes deserve further scrutiny for their potential roles in cancer immunotherapy, particularly in renal cell carcinomas.

## DISCUSSION

The domain of immune checkpoint (IC) expression has been largely untouched in canine cancers. Here, we provide a comprehensive overview of IC expression patterns in both human and canine cancers. We unveil potential canine models for human cancer immunotherapy and identify prospective IC targets that intersect both species. By comparing the ‘IC signatures’ encompassing 44 IC genes in 27 human and 14 canine cancer types, we uncover distinct and shared patterns across species, cancer types, and individual patients. Our findings advance the fields of human cancer immunology and canine immunology and fill a knowledge gap in comparative oncology.

### Prior Studies

The veterinary part of our investigation builds upon prior studies that identified PD-1 receptors on canine T cells and observed PD-L1 in various canine cancer types using immunohistochemistry^51–54^. Other authors reported the presence of CTLA-4 and IDO in specific canine cancer types, such as B-cell lymphoma and melanoma^55, 56^, and found that B7-H4 was overexpressed in bladder cancer^54^. Murakami *et al*. advanced the field with their RT‒qPCR investigation of 20 immune-regulating molecules in fresh samples^57^, yet their study was constrained by a small sample size covering a wide range of cancer subtypes. Our study expands the scope and depth of the investigation with a larger sample size and comprehensive IC profiling.

### The Immune Environment of Canine Cancers

Our study reveals the expression of 44 ICs in the majority of the 14 analyzed canine cancers. The abundance of these genes varied across cancer types, highlighting the potential for both universal and cancer type-specific treatment targets. Additionally, we observed diverse levels of immune cell infiltration and varied levels of estimated immune inhibition, thus characterizing the tumor microenvironment in aspects crucial for successful immunotherapy.

### Immune Checkpoint Signatures That Link Human and Dog Cancers

We used hierarchical clustering to compare IC signatures between canine and human cancers. This allowed us to discover remarkable similarities between canine glioma and human glioma, as well as between canine osteosarcoma and human sarcoma, highlighting them as potential models for immunotherapy development. Intriguingly, this analysis also suggested a close relationship between brain cancers, human melanoma and human renal cell carcinoma in the IC context. Given the successful treatment of melanoma with immune checkpoint blockade (ICB), this association could provide some rationale for exploring ICB therapies for brain cancers. Furthermore, using PCA, we found unexpected similarity between human chronic lymphocytic leukemia cases and canine malignancies at large. Further research is needed to validate these findings.

### Brain Cancers, Immunotherapy and Comparative Oncology

The striking similarities in immune checkpoint expression human and canine glioma were evidenced through UMAP, PCA, and hierarchical clustering analysis. In fact, this similarity was unparalleled by any other pair of cancers in this study, including intraspecies pairs. Canine gliomas also exhibited high immune inhibition scores, increased T-reg infiltration, and exhausted lymphocytes, mirroring the immunologically ‘cold’ nature of human gliomas. One notable finding was the significant expression of SIRPA, an inhibitory receptor on macrophages, and its ligand CD47, which is often overexpressed in cancers. SIRPA is also prevalent in healthy brain tissues; nevertheless, its high level in brain cancers suggests that the SIRPA-CD47 interaction may be highly important in gliomas across both species. By highlighting the shared IC landscape in human and canine gliomas, our findings lend further support to the suitability of canine gliomas as models for human gliomas and point to SIRPA as a promising target for glioma immunotherapy. Although many researchers have doubted the feasibility of immunotherapy for glioma, this challenging cancer offers an opportunity to explore key innovative treatment approaches: novel animal models, personalized therapies, combination treatments, and targeting of immune populations beyond T-cells. Although we observed close similarity between canine glioma and human meningioma - another brain cancer - we prioritized gliomas and glioblastomas due to their aggressive nature and lower operability. Insights from our glioma analysis may extend to meningiomas. For an in-depth discussion on glioma, please see the supplementary materials.

### NKG2A vs NK Cells in Canine Glioma

We observed an apparent anomaly in canine glioma infiltration. Unlike other canine cancers, glioma presented detectable expression of NKG2A, a known receptor of NK cells. However, it appeared to simultaneously present with the lowest expression of NK marker genes of all analyzed cancers. Although we have no definitive explanation for this phenomenon, we propose a hypothesis based on alternative NKG2A roles in humans. It has been reported that NKG2A can be expressed by Natural Killer T (NKT) cells, a unique group of T lymphocytes that possess characteristics of both T cells and NK cells. Furthermore, a subset of CD8+ αβ T-cells, particularly γδ T-cells, is also known to express NKG2A^58, 59^. NKG2A is additionally present in some T-cells that infiltrate peripheral nerves^60^. Therefore, the unexpected relationship between comparatively high NKG2A abundance and low NK cell score in canine glioma could be attributed to the involvement of noncanonical NKG2A carrier cells. This, however, requires dedicated research to prove.

### Notes on Osteosarcoma

Osteosarcoma may be unlikely to benefit from targeting inhibitory ICs like CTLA-4, PD-1 and OX40 alone^61^. Based on our canine IC expression heatmap and human literature, we suggest that B7-H3, a complex-function IC molecule associated with poor prognosis^62^ that is highly expressed in both human and canine OS, could be an interesting target for studying the human-canine intersection.

### Clustered IC Signatures

In hierarchical clustering analysis, most cancer signatures aggregated based on species, cancer histological type and primary site. These findings shed light on the links between cancer characteristics and the immune checkpoint landscape. Within human cancer subtypes, further research should explore the underlying molecular mechanisms driving these trends, particularly in regard to optimizing treatment. By exploring species-specific IC differences, we identified four functionally unrelated IC genes (CD160, A2AR, NKG2A, and OX40) that drove the divide between canine and human IC signatures in PCA. Since the human and canine datasets were obtained from different sources, further research needs to validate these findings. However, as most IC genes did not exhibit statistically significant differential expression between species, bias is an unlikely explanation.

### IC Signatures of Healthy and Malignant Tissues

In this investigation, we utilized UMAP and PCA to compare the IC signatures of cancer tissues and corresponding healthy tissues in a few human cancer types. In UMAP, renal cell carcinoma and chromophobe renal cell carcinoma exhibited a distinct shift between healthy and diseased states. In addition, we observed a significant contrast between consistent signatures of healthy lung samples and highly diverse signatures of adenocarcinoma cases. These results suggest that the transition of healthy tissue into a cancerous one involves a broad shift in immune checkpoint expression. The diversity observed in lung adenocarcinoma sheds light on the heterogeneity of IC-based immune evasion strategies within one cancer type. On the other hand, these trends were not clear in cholangiocarcinoma and hepatocarcinoma, but low numbers of healthy reference samples limited interpretability in these cases. PCA mirrored the above patterns, especially in renal carcinomas. In renal cancers, the main contributors to PC-1, the axis separating healthy and sick – were the CD86, TIGIT, ICOS, and BTLA genes. Further exploration could focus on their role in cancer development and potential as immunotherapy targets. Importantly, healthy reference samples are typically derived from tumor-adjacent tissues, which might share certain features with nearby cancer tissues. Consequently, this effect may downplay the real differences between healthy and cancerous tissues in this experimental setup.

### The IC Signatures of Individual Patients

Our analysis revealed that human patients with the same cancer type tended to exhibit similar IC signatures. For example, we noted a close alignment of IC signatures among glioma cases within species, which is indicative of conserved immune characteristics. However, in some cancers, such as sarcoma, the case signatures were more heterogeneous, hinting at greater variability. Cancers displaying uniform IC signatures across patients offer promise for more consistent treatment success with a single therapeutic strategy. In contrast, the observation of cancers with high signature variability underscores the need for personalized therapy based on the unique immune landscape of each patient. Therefore, our comprehensive characterization of IC signatures complements individualized therapeutic approaches such as cancer immunogram^63^. Further investigation is needed to understand the factors underlying differences in IC signatures and their implications for treatment outcomes.

### Differential Gene Expression Between Identified Clusters

When we calculated differential IC gene expression between the previously defined clusters of human cancers, many trends were observed, which requires further validation. Interestingly, B7-H4 was characteristically upregulated in female cancers, consistent with studies of ovarian, cervical, and breast cancers^64–69^. Gynecological adenocarcinomas appeared to differentiate themselves from other adenocarcinomas via increased IDO expression. These findings showcase both the shared traits and unique IC characteristics of gynecological adenocarcinomas.

The results of canine-human clusters comparison by differential expression and PCA did not converge, except for A2AR, highlighting it as the most confident species-related gene. In humans, A2AR can be upregulated in response to anti-PD-1 treatment, and its blockade increases treatment efficacy^70^. While we observed trends in cancer type clusters, the heterogeneity of the IC signature across individual patients with the same diagnosis appears to be a crucial research direction to explore.

### Cross-species target conservation

As we assessed the conservation of IC amino acid sequences between dog and human (**Tab. S6**), TIM-3 was the least conserved (24/100), and B7-H4 together with NECTIN-4 were the most conserved (94/100) of the analyzed proteins. While these scores do not specifically describe extracellular domains containing potential epitopes, some canine ICs may be druggable with caninized human antibodies. Others will likely require the development of canine-specific antibodies. Although the cross-reactivity of human antibody therapeutics is disputed, Pantelyushin *et al*. convincingly described ipilimumab, nivolumab, atezolizumab and avelumab (which target CTLA-4, PD-1 and PD-L1, respectively) as cross-reactive, with various levels of functionality^71^. In this context, Nectin-4 is a particularly interesting target, combining high conservation, IC inhibition and tumor specificity^72^.

### Limitations

The limitations of our study were primarily dictated by the available data. Although the selected human dataset did not include osteosarcoma, it included 27 other precisely diagnosed cancer types, including some closely related ones. In contrast, we could only analyze 14 distinct types of canine cancer. This discrepancy is due to a paucity of canine studies and the less precise diagnostic categorization of canine cancers. We anticipate that future subdivisions of canine cancers into discrete subtypes will allow for more precise comparisons with human equivalents, thus revealing greater interspecies similarity. This study focused on 44 ICs, but many have likely not yet been identified, which precludes capturing the full complexity of immune checkpoint landscapes. Furthermore, we could not analyze several ICs due to their apparent absence in the CanFam6 reference transcriptome; these included SIGLEC7/9, SLAMF3/4, LILRB1-5, BTN2A1, and KIRs. Multiple gene markers of immune cell populations were also excluded due to their absence or misrepresentation in the reference transcriptome (see supplementary methods). These research limitations will diminish as the canine model becomes more widespread.

A common challenge in transcriptomic and proteomic studies of solid tumors, including ours, is identifying the cell type responsible for the presence of the analyte. IC ligands, in particular, can be expressed by multiple cell types within the tumor microenvironment. However, the source of ICs doesn’t significantly impact our evaluation of the IC landscape. Regardless of the source, inhibitory ligands interact with immune effector cells and are potential ICB targets. One caveat is the possibility that cancer cells themselves express functional IC receptors, a hypothesis lacking consistent evidence. We followed the common assumption that comparatively high IC expression indicates a promising target.

The functional equivalence of ICs in canines and humans is not guaranteed. Interspecies variations in IC functionality, as seen in GITR between mice and humans, may bolster the relevance of the canine model but warrant caution in studying isolated ICs^73^. Moreover, genetic and metabolic differences between the two species may limit the applicability of the findings to humans. Despite promising results, it is essential to validate the functional equivalence of human and canine ICs.

Transcript abundance, as measured by RNAseq, does not perfectly correlate with the production and membrane presence of the associated protein^74–77^. However, transcriptomics provides a snapshot of the cell’s signaling frozen in time, offering invaluable insights. Flow cytometry and immunohistochemistry can help validate our findings as more antibodies against canine ICs become available.

### Future Perspectives

One important aspect of the immune checkpoint signatures that remained beyond the scope of this study is their interplay with therapy response, as well as the emergence of resistance, especially via the upregulation of undrugged ICs. Additionally, the possibility of predicting hyperprogressive disease based on these signatures is a promising question. In light of the increasing development costs of novel immunotherapeutics and challenges in thoroughly evaluating their effects before approval, streamlining preclinical drug screening and development is critical. Here, we revealed key similarities in canine and human immune checkpoint expression patterns in cancer, thereby reinforcing the argument for comparative oncology. Our analysis of individual human cancer IC signatures can pave the way for more personalized treatment decisions and monitoring. Ultimately, we hope these insights will catalyze advancements in both veterinary and human oncology, leading to the generation of more accessible, effective, and safer immunotherapies.

## Acknowledgments

We would like to thank the authors of all the cited canine studies as well as the PCAWG consortium for openly sharing and describing the datasets, thus enabling this work. Additionally, we thank the CI-TASK, Gdansk and PLGrid Infrastructure, Poland, for providing computational resources. MK acknowledges that portions of this work have been derived from a chapter of his doctoral thesis, which is currently under review and not publicly accessible as of the date of this manuscript. The thesis, titled “Of Dogs and Men. Tracing Immune Checkpoint Signatures Across Cancers and Unleashing the Potential of Canine PD-1 Antibodies”, was submitted to the University of Gdansk.

This work was performed at the International Centre for Cancer Vaccine Science, a project carried out within the International Research Agendas programme of the Foundation for Polish Science co-financed by the European Union under the European Regional Development Fund [grant MAB/3/2017].

## Author Contributions

**M. Kocikowski** - conceptualization, data curation, formal analysis, investigation, methodology, project administration, software, visualization, writing–original draft, writing– review and editing. **M. Yébenes Mayordomo** - methodology, software, writing–review and editing. **J.A. Alfaro** - conceptualization, writing–review and editing, resources, supervision. **M. Parys** - conceptualization, writing–review and editing, resources, supervision.

## Conflicts of Interest

MP is a founder of CanCan Diagnostics, which had no involvement in the study and its results, and declares no conflict of interest. The remaining authors declare that they have no conflicts of interest.

## Data Availability

Publicly available datasets were analyzed in this study. These data can be found under the IDs displayed in **Table S5**.

