## Supplemental File for "Barking Up the Right Tree: Immune Checkpoint Signatures of Human and Dog Cancers"

### Supplementary methods

#### Marker gene choice

The CPA3 (mast cell carboxypeptidase A) gene was removed from the results as misleading. We observed an uncharacteristically high CPA3 apparent abundance in insulinoma (a pancreatic cancer), unlikely to originate from mast cells, while pancreatic carboxypeptidase B (CPB2) - a CPA3 paralogue gene - did not exhibit similarly high expression. We performed an Ensembl blast of the canonical CPA3 transcript (ENSCAFT00000071516.2) cDNA sequence against cDNA (transcripts/splice variants) of all Ensembl dog breeds (German Shepherd, ROS_Cfam_1.0, Basenji_breed-1.1. Great Dane and the Boxer itself). In all breeds except the Boxer, the top hits belonged to CPB2. Additionally, we observed similarly high expression of CPA1 and CPA5 in insulinoma. Suspecting a confusion in the reference genome/transcriptome annotation we contacted the Ensembl support. They have kindly checked and confirmed the issue stems from inaccurate annotation of the locus encoding for both CPA3 and CPB2 in the Ensembl dog breeds. In our case readthrough transcripts (ENSCAFT00000071516.2 and ENSCAFT00000087051.2) reportedly joined both transcript sets because of overlapping protein-coding regions. Until the locus is reviewed, we resigned from analyzing CPA3 and CPB expression, which would require modification of the reference used at the fastq files quantification stage.

#### Supplementary Discussion

##### Glioma vs immunotherapy

The parallels between human and canine glioma (and to a large extent canine meningioma and human glioblastoma) necessitate a deeper discussion. The IC signatures of these cancers of the two species presented remarkable similarity in 3 different analyses: UMAP, PCA, and hierarchical clustering of distances. This we consider a very strong evidence for the canine glioma resemblance of human glioma in terms of immune checkpoint expression.

Consistent with the rather immunologically ‘cold’ character of human gliomas, in canine gliomas we observed a high immune inhibition score, as well as high score for T-reg cells and exhausted lymphocytes. The IC expression was relatively low, barring the notably abundant SIRPA - an inhibitory receptor found predominantly on macrophages. Its ligand, CD47 - a broadly expressed membrane protein often overexpressed by cancers - was present at a considerable level.

Our IC-focused findings support canine gliomas as a suitable model for human gliomas, which has in the past been backed on the level of genomic events, methylation and immune infiltrate[1]. In the cited study, canine gliomas appeared to recapitulate human pediatric gliomas particularly well. In our analysis of the IC landscape, the similarity in gliomas of both species was pronounced despite the fact that the median patient age was 40.5 years in the glioma dataset we used (calculated based on Table S1 from[2] and patient IDs from EBI PCAWG experiment design table). The human-canine glioma similarity has been exploited in pilot drug studies[3–6] and precipitated more clinical assessments[6]. One team investigated peptide-based inhibition of a CD200 purported IC in high-grade canine gliomas with promising outcomes[7]. However, antibody-based blockade or the major checkpoints has not been studied in the canine model yet.

The lack of such studies can likely be traced to the limited success of the ICB interventions in human glioma and glioblastoma. However, one could argue these unsatisfactory results only call for more investment in innovative research approaches. Moreover, new studies begin to challenge the notion of glioma as untreatable with immunotherapy[8–10].

Still, there is no shortage of challenges in studying and treating gliomas, unique as they are. These issues have been elegantly set in the wider theme of cancer immunology, and comprehensively synthesized, by Khasraw and colleagues[11]. To name a few, the availability and volume of the samples is limited and repeated biopsies are not possible due to the tumor location. These factors limit the statistical power of studies and the range of assays that can be run. The unique brain environment, with its specific immune features, hypoxia, and other unusual physiological conditions, is particularly difficult to mimic realistically in animal models, both syngeneic and in xenografts. What’s more, the knowns established in other cancers, become unknowns in glioma. For instance, tumor mutational burden (TMB), commonly considered a proxy for tumor immunogenicity and a predictor of ICB success, does not hold the same predictive power in glioma as in other tumors[12,13]. The limited understanding of glioma led the field to try treatments that were promising in other cancers rather than based on glioma-specific rationale. Researchers do call for novel preclinical animal models for studying glioma and glioblastoma[11]. This is where the canine model has a special role to play.

Yet another troublesome characteristic of gliomas is the myeloid character of infiltrating immune cells, which discourages the PD-1 ICB and similar therapeutic approaches that are theoretically aimed at T-cells. It is worthwhile mentioning that the theoretical assumptions of ICB do not cover all its effects - it is becoming clear that B, NK and other immune cells meaningfully react to those treatments and contribute to its success or failure[14,15]. More importantly, the prevalence of immune cells of myeloid lineage - such as macrophages, neutrophils and dendritic cells - in gliomas may be the key to treat these cancers successfully.

SIRPA, the inhibitory IC receptor that we detected on a comparatively high level in canine glioma, is a critical mediator of immune responses in myeloid cells. Its interaction with CD47 plays a significant role in cancer immune evasion. SIRPA is naturally highly expressed in the brain[16]. However, elevated levels of SIRPA and CD47 correlate with decreased survival in human glioma and glioblastoma[17]. The CD47/SIRPA interaction in glioblastoma broadly inhibits immune cells, rendering ICB a promising treatment strategy[18,19].

Targeting SIRPA - rather than its ligand CD47 - may be beneficial, reaching the immune cells of interest similarly, rather than the wide array of cells expressing CD47. Targeting SIRPA has been made difficult by SIRPA's low conservation between species (**Tab. 4** - Methods) and high polymorphism within the CD47-binding domain. However, one team has developed pan-allelic, 'pan-mammal' antibodies targeting human, monkey and mouse SIRPA and blocking its interaction with CD47[20,21]. The team has utilized another unusual animal model for expanding the capabilities of cancer research. Uniquely, they raised the antibodies in chickens. The high phylogenetic distance between chickens and humans, together with low homology of their SIRPA sequences, became a captured opportunity in this case. These chicken traits allowed for raising antibodies against unique and pan-mammalian epitopes that would not be accessible, were the antibodies raised in mice or rabbits[21]. This is to say that antibodies raised in mice would likely not recognize epitopes of murine SIRPA or many similar mammalian epitopes. While these researchers did not aim at antibodies cross-reactive with canine SIRPA, and did not seem to evaluate their activity against the canine protein variant, their elegant approach seems to hold promise of developing such antibodies.

Despite many unique opportunities for immunotherapy development, many still believe glioma and glioblastoma are not treatable with such modality. We propose the high-grade glioma is both a cancer of high unmet need, and a chance for cancer immunology to progress through the application of novel animal models. It is also fertile ground for testing key paradigm changes, such as targeting of checkpoints beyond PD-1, personalized therapy recognizing tumor heterogeneity, combinatorial treatments involving appropriate methods for their evaluation, and targeting all of the relevant immune cell populations.

#### Supplementary tables

**Table S1: Mean-based differential IC expression comparing human and canine cancer types and clusters.** Genes that were found significant in both median- and mean-based analysis were marked in bold for denoting increased confidence. Fc - fold change, log2fc - Log of fc, pval - p value, HCC - hepatocellular carcinoma, MEL - melanoma, RCC - renal cell carcinoma, SCC - squamous cell carcinoma, TCC - transitional cell carcinoma.

| Comparison | gene_id | mean TPM | reference TPM | fc | log2_fc | zscore | pval | regulation |
|---|---|---|---|---|---|---|---|---|
| Canine vs Human | PVRIG | 11.07 | 0.54 | 20.33 | 4.35 | 2.09 | 1.82E-02 | up |
| | **A2AR** | 8.96 | 0.51 | 17.40 | 4.12 | 1.98 | 2.38E-02 | up |
| Sarcoma vs other (human) | **B7-H4** | 0.10 | 33.92 | 0.00 | -8.41 | -2.94 | 1.64E-03 | down |
| | FGL-1 | 0.10 | 24.58 | 0.00 | -7.94 | -2.74 | 3.11E-03 | down |
| | **NECTIN4** | 0.40 | 43.96 | 0.01 | -6.78 | -2.23 | 1.30E-02 | down |
| | Arginase 1 | 0.30 | 14.84 | 0.02 | -5.63 | -1.72 | 4.28E-02 | down |
| | BTLA | 0.20 | 9.07 | 0.02 | -5.50 | -1.66 | 4.81E-02 | down |
| Melanoma vs other (human) | **B7-H4** | 0.10 | 33.92 | 0.00 | -8.41 | -3.65 | 1.33E-04 | down |
| | **NECTIN4** | 0.50 | 43.96 | 0.01 | -6.46 | -2.62 | 4.35E-03 | down |
| | Arginase 1 | 0.20 | 14.84 | 0.01 | -6.21 | -2.49 | 6.30E-03 | down |
| HCC vs other (human) | **NECTIN4** | 0.20 | 43.96 | 0.00 | -7.78 | -2.66 | 3.86E-03 | down |
| | B7-H4 | 0.30 | 33.92 | 0.01 | -6.82 | -2.24 | 1.24E-02 | down |
| | BTLA | 0.20 | 9.07 | 0.02 | -5.50 | -1.67 | 4.79E-02 | down |
| | **Arginase 1** | 335.00 | 14.84 | 22.58 | 4.50 | 2.72 | 3.26E-03 | up |
| | **FGL-1** | 530.00 | 24.58 | 21.57 | 4.43 | 2.69 | 3.56E-03 | up |
| | LIGHT | 16.00 | 3.01 | 5.31 | 2.41 | 1.80 | 3.56E-02 | up |
| Gynecological cancers vs other (human) | FGL-1 | 0.08 | 24.58 | 0.00 | -8.20 | -4.28 | 9.55E-06 | down |
| | Arginase 1 | 0.22 | 14.84 | 0.01 | -6.10 | -3.07 | 1.06E-03 | down |
| Gastric cluster vs other (human) | BTLA | 0.30 | 9.07 | 0.03 | -4.92 | -3.28 | 5.13E-04 | down |
| | **B7-H4** | 3.36 | 33.92 | 0.10 | -3.34 | -1.90 | 2.86E-02 | down |
| | Arginase 1 | 1.50 | 14.84 | 0.10 | -3.31 | -1.88 | 3.03E-02 | down |
| | **CD70** | 1.34 | 13.00 | 0.10 | -3.28 | -1.85 | 3.20E-02 | down |
| | CD155 | 68.40 | 34.15 | 2.00 | 1.00 | 1.88 | 2.97E-02 | up |
| | LIGHT | 5.80 | 3.01 | 1.93 | 0.95 | 1.84 | 3.32E-02 | up |
| Lymphomas vs other (human) | **NECTIN4** | 0.50 | 43.96 | 0.01 | -6.46 | -3.54 | 2.00E-04 | down |
| | Arginase 1 | 0.35 | 14.84 | 0.02 | -5.41 | -3.04 | 1.18E-03 | down |
| | **B7-H3** | 7.00 | 56.71 | 0.12 | -3.02 | -1.91 | 2.81E-02 | down |
| | **CD155** | 4.50 | 34.15 | 0.13 | -2.92 | -1.87 | 3.11E-02 | down |
| Human brain cancers vs other human | **NECTIN4** | 0.15 | 43.96 | 0.00 | -8.20 | -2.22 | 1.33E-02 | down |
| | B7-H4 | 0.30 | 33.92 | 0.01 | -6.82 | -1.67 | 4.78E-02 | down |
| | SIRPA | 180.00 | 60.75 | 2.96 | 1.57 | 1.70 | 4.48E-02 | up |
| Brain cancers with melanoma and RCC vs other (human) | **NECTIN4** | 0.25 | 43.96 | 0.01 | -7.46 | -2.78 | 2.69E-03 | down |
| | FGL-1 | 0.15 | 24.58 | 0.01 | -7.36 | -2.73 | 3.12E-03 | down |
| | B7-H4 | 0.30 | 33.92 | 0.01 | -6.82 | -2.48 | 6.61E-03 | down |
| | Arginase 1 | 0.25 | 14.84 | 0.02 | -5.89 | -2.03 | 2.11E-02 | down |
| SCC and TCC vs other (human) | FGL-1 | 0.03 | 24.58 | 0.00 | -9.94 | -4.67 | 1.52E-06 | down |
| | Arginase 1 | 0.20 | 14.84 | 0.01 | -6.21 | -2.70 | 3.51E-03 | down |
| | BTLA | 0.43 | 9.07 | 0.05 | -4.42 | -1.75 | 4.05E-02 | down |
| Carcinomas vs other (human) | BTLA | 0.55 | 9.07 | 0.06 | -4.06 | -3.97 | 3.65E-05 | down |
| | CTLA-4 | 2.44 | 17.79 | 0.14 | -2.87 | -2.61 | 4.54E-03 | down |
| | CD27 | 6.34 | 26.73 | 0.24 | -2.08 | -1.70 | 4.42E-02 | down |
| Adenocarcinomas vs other (human) | Arginase 1 | 0.22 | 14.84 | 0.01 | -6.08 | -3.55 | 1.93E-04 | down |
| | FGL-1 | 0.40 | 24.58 | 0.02 | -5.94 | -3.46 | 2.71E-04 | down |
| | BTLA | 0.59 | 9.07 | 0.07 | -3.94 | -2.13 | 1.68E-02 | down |
| Gynecological adenocarcinomas vs other adenocarcinomas (human) | GITRL | 0.35 | 1.04 | 0.34 | -1.58 | -2.61 | 4.54E-03 | down |
| Gastric adenocarcinomas vs other adenocarcinomas (human) | **B7-H4** | 2.53 | 93.29 | 0.03 | -5.20 | -4.01 | 3.09E-05 | down |
| | CD70 | 2.00 | 21.54 | 0.09 | -3.43 | -2.48 | 6.49E-03 | down |

**Table S2: Median abundance of the four species-dividing genes quantified in TPM units;** values provided only as an illustration of the general trends.

| | **CD160** | **A2AR** | **NKG2A** | **OX40** |
|---|---|---|---|---|
| **MEDIAN DOG TPM:** | 2.29 | 4.27 | 0.00 | 0.87 |
| **MEDIAN HUMAN TPM:** | 0.30 | 0.20 | 0.40 | 6.00 |

**Table S3: The numbers of malignant and healthy samples available in the analyzed human cancers.**

| Human cancer type | Cancer samples | Normal samples |
|---|---|---|
| B-cell non-Hodgkin lymphoma | 103 | 0 |
| bladder transitional cell carcinoma | 23 | 4 |
| breast adenocarcinoma | 85 | 6 |
| cervical adenocarcinoma | 2 | 0 |
| cervical squamous cell carcinoma | 18 | 0 |
| cholangiocarcinoma | 18 | 16 |
| chromophobe renal cell carcinoma | 43 | 14 |
| chronic lymphocytic leukemia | 68 | 0 |
| colorectal adenocarcinoma | 51 | 0 |
| endometrial adenocarcinoma | 44 | 1 |
| esophageal adenocarcinoma | 7 | 0 |
| follicular thyroid carcinoma | 47 | 4 |
| gastric adenocarcinoma | 29 | 2 |
| glioblastoma multiforme | 28 | 0 |
| glioma | 18 | 0 |
| head and neck squamous cell carcinoma | 42 | 1 |
| hepatocellular carcinoma | 100 | 53 |
| invasive lobular carcinoma | 6 | 0 |
| lung adenocarcinoma | 37 | 9 |
| lymphoma | 2 | 0 |
| melanoma | 36 | 0 |
| ovarian adenocarcinoma | 101 | 0 |
| pancreatic adenocarcinoma | 75 | 0 |
| prostate adenocarcinoma | 19 | 1 |
| renal cell carcinoma | 117 | 37 |
| sarcoma | 34 | 0 |
| squamous cell lung carcinoma | 47 | 2 |
| **TOTAL:** | **1200** | **150** |

**Table S4: The distance matrix compares the immune checkpoint signatures between human and canine cancer types;** the values are a measure of distance, hence lower numbers mean higher similarity between two cancer types.

| | ameloblastoma (canine) | B-cell lymphoma (canine) | glioma (canine) | hemangiosarcoma (canine) | insulinoma (canine) | lung carcinoma (canine) | mammary cancers (canine) | melanoma (canine) | meningioma (canine) | oral squamous cell carcinoma (canine) | osteosarcoma (canine) | prostate carcinoma (canine) | T-cell lymphoma (canine) | urothelial carcinoma (canine) |
|---|---|---|---|---|---|---|---|---|---|---|---|---|---|---|
| ameloblastoma (canine) | | | | | | | | | | | | | | |
| B cell lymphoma (canine) | 0.383 | | | | | | | | | | | | | |
| glioma (canine) | 0.376 | 0.728 | | | | | | | | | | | | |
| hemangiosarcoma (canine) | 0.141 | 0.355 | 0.275 | | | | | | | | | | | |
| insulinoma (canine) | 0.124 | 0.373 | 0.405 | 0.173 | | | | | | | | | | |
| lung carcinoma (canine) | 0.188 | 0.463 | 0.352 | 0.275 | 0.161 | | | | | | | | | |
| mammary cancers (canine) | 0.142 | 0.453 | 0.275 | 0.255 | 0.209 | 0.174 | | | | | | | | |
| melanoma (canine) | 0.291 | 0.551 | 0.333 | 0.304 | 0.199 | 0.208 | 0.266 | | | | | | | |
| meningioma (canine) | 0.327 | 0.724 | **0.083** | 0.340 | 0.301 | 0.196 | 0.240 | 0.244 | | | | | | |
| oral squamous cell carcinoma (canine) | 0.222 | 0.404 | 0.565 | 0.376 | 0.324 | 0.416 | 0.347 | 0.493 | 0.496 | | | | | |
| osteosarcoma (canine) | 0.293 | 0.675 | 0.438 | 0.333 | 0.242 | 0.296 | 0.378 | 0.302 | 0.328 | 0.554 | | | | |
| prostate carcinoma (canine) | 0.163 | 0.452 | 0.199 | 0.236 | 0.266 | 0.266 | 0.118 | 0.270 | 0.242 | 0.344 | 0.400 | | | |
| T cell lymphoma (canine) | 0.778 | 0.755 | 0.949 | 0.821 | 0.748 | 0.747 | 0.770 | 0.739 | 0.857 | 0.899 | 0.871 | 0.887 | | |
| urothelial carcinoma (canine) | 0.248 | 0.520 | 0.609 | 0.500 | 0.434 | 0.361 | 0.174 | 0.545 | 0.556 | 0.336 | 0.609 | 0.232 | 0.927 | |
| B cell non Hodgkin lymphoma (human) | 0.814 | 0.571 | 0.784 | 0.683 | 0.804 | 0.609 | 0.832 | 0.742 | 0.779 | 0.914 | 0.862 | 0.765 | 0.832 | 0.906 |
| chronic lymphocytic leukemia (human) | 0.697 | 0.617 | 0.927 | 0.781 | 0.645 | 0.628 | 0.748 | 0.680 | 0.801 | 0.819 | 0.751 | 0.816 | 0.832 | 0.848 |
| lymphoma (human) | 0.939 | 0.690 | 0.837 | 0.802 | 0.944 | 0.734 | 0.950 | 0.881 | 0.857 | 1.017 | 0.967 | 0.882 | 0.846 | 1.015 |
| glioblastoma multiforme (human) | **0.416** | 0.745 | **0.070** | **0.301** | **0.426** | **0.381** | **0.343** | **0.334** | **0.123** | 0.621 | **0.268** | **0.248** | 0.966 | 0.657 |
| glioma (human) | **0.497** | 0.882 | **0.025** | **0.416** | **0.523** | **0.441** | **0.358** | **0.394** | **0.099** | 0.682 | **0.466** | **0.266** | 0.978 | 0.696 |
| breast adenocarcinoma (human) | **0.473** | 0.843 | 0.639 | 0.644 | 0.641 | **0.505** | **0.356** | 0.619 | 0.607 | 0.637 | **0.391** | **0.399** | 1.060 | **0.245** |
| invasive lobular carcinoma (human) | **0.447** | 0.801 | 0.655 | 0.655 | 0.648 | **0.486** | **0.306** | 0.659 | 0.620 | 0.594 | **0.522** | **0.383** | 1.038 | **0.150** |
| esophageal adenocarcinoma (human) | **0.287** | 0.569 | 0.568 | **0.331** | **0.365** | **0.417** | **0.388** | **0.345** | 0.567 | **0.500** | **0.341** | **0.303** | 0.914 | **0.418** |
| chromophobe renal cell carcinoma (human) | 0.702 | 0.820 | 0.794 | 0.633 | 0.627 | 0.622 | 0.745 | **0.480** | 0.737 | 0.829 | **0.489** | 0.699 | 0.976 | 0.828 |
| renal cell carcinoma (human) | 0.552 | 0.704 | **0.175** | **0.384** | **0.510** | **0.499** | **0.429** | **0.295** | **0.245** | 0.744 | **0.492** | **0.306** | 0.961 | 0.740 |
| colorectal adenocarcinoma (human) | **0.321** | **0.419** | 0.631 | **0.311** | **0.355** | **0.471** | **0.442** | **0.387** | 0.661 | **0.475** | **0.328** | **0.360** | 0.889 | **0.506** |
| cholangiocarcinoma (human) | **0.478** | 0.639 | **0.450** | **0.421** | **0.357** | **0.465** | **0.392** | **0.230** | **0.473** | 0.662 | **0.412** | **0.386** | 0.928 | 0.641 |
| hepatocellular carcinoma (human) | 1.006 | 1.064 | 0.951 | 1.021 | 0.874 | 1.022 | 0.969 | 1.015 | 0.975 | 0.930 | 1.014 | 1.049 | 1.144 | 1.073 |
| lung adenocarcinoma (human) | **0.181** | **0.451** | **0.498** | **0.314** | **0.257** | **0.190** | **0.280** | **0.324** | **0.384** | **0.398** | **0.225** | **0.260** | 0.835 | **0.326** |
| squamous cell lung carcinoma (human) | **0.393** | 0.757 | 0.607 | **0.532** | 0.577 | 0.556 | **0.441** | 0.572 | 0.590 | 0.559 | **0.322** | **0.368** | 0.996 | **0.384** |
| head and neck squamous cell carcinoma (human) | **0.276** | 0.777 | **0.525** | **0.469** | **0.498** | **0.498** | **0.373** | **0.530** | **0.509** | **0.496** | **0.371** | **0.277** | 0.958 | **0.327** |
| ovarian adenocarcinoma (human) | **0.398** | 0.666 | 0.669 | 0.559 | **0.417** | **0.233** | **0.236** | **0.504** | **0.517** | 0.590 | **0.433** | **0.448** | 0.892 | **0.237** |
| pancreatic adenocarcinoma (human) | **0.262** | 0.594 | **0.500** | **0.380** | **0.394** | **0.213** | **0.357** | **0.394** | **0.403** | **0.490** | **0.339** | **0.317** | 0.890 | **0.346** |
| prostate adenocarcinoma (human) | **0.335** | 0.764 | 0.599 | **0.464** | **0.360** | **0.343** | **0.420** | **0.318** | **0.474** | 0.602 | **0.154** | **0.415** | 0.907 | **0.485** |
| sarcoma (human) | **0.454** | 0.794 | **0.444** | **0.369** | **0.432** | **0.491** | 0.545 | **0.420** | **0.448** | 0.723 | **0.086** | **0.464** | 0.985 | 0.759 |
| melanoma (human) | **0.520** | 0.818 | **0.244** | **0.437** | **0.467** | **0.476** | **0.445** | **0.239** | **0.278** | 0.731 | **0.219** | **0.328** | 0.966 | 0.737 |
| gastric adenocarcinoma (human) | **0.264** | **0.362** | **0.380** | **0.183** | **0.305** | **0.344** | **0.341** | **0.361** | **0.439** | **0.430** | **0.303** | **0.233** | 0.905 | **0.491** |
| follicular thyroid carcinoma (human) | **0.407** | 0.658 | 0.604 | **0.392** | **0.358** | **0.368** | **0.498** | **0.385** | **0.494** | 0.673 | **0.287** | **0.527** | 0.847 | 0.685 |
| bladder transitional cell carcinoma (human) | 0.558 | 0.946 | 0.909 | 0.855 | 0.852 | 0.856 | 0.659 | 0.833 | 0.899 | 0.633 | 0.759 | 0.571 | 1.084 | **0.373** |
| cervical adenocarcinoma (human) | 0.668 | 0.858 | 0.888 | 0.787 | 0.685 | 0.654 | 0.659 | 0.741 | 0.819 | 0.833 | 0.694 | 0.626 | 1.001 | 0.590 |
| cervical squamous cell carcinoma (human) | **0.360** | 0.785 | 0.673 | 0.563 | 0.576 | 0.575 | **0.436** | 0.591 | 0.646 | 0.544 | **0.414** | **0.375** | 0.955 | **0.344** |
| endometrial adenocarcinoma (human) | **0.482** | 0.656 | 0.747 | 0.593 | 0.554 | **0.520** | **0.312** | 0.634 | 0.737 | 0.615 | **0.459** | **0.421** | 0.990 | **0.240** |

| | B-cell non-Hodgkin lymphoma (human) | chronic lymphocytic leukemia (human) | lymphoma (human) | glioblastoma multiforme (human) | glioma (human) | breast adenocarcinoma (human) | invasive lobular carcinoma (human) | esophageal adenocarcinoma (human) | chromophobe renal cell carcinoma (human) | renal cell carcinoma (human) | colorectal adenocarcinoma (human) | cholangiocarcinoma (human) | hepatocellular carcinoma (human) | lung adenocarcinoma (human) | squamous cell lung carcinoma (human) | head and neck squamous cell carcinoma (human) | ovarian adenocarcinoma (human) | pancreatic adenocarcinoma (human) | prostate adenocarcinoma (human) | sarcoma (human) | melanoma (human) | gastric adenocarcinoma (human) | follicular thyroid carcinoma (human) | bladder transitional cell carcinoma (human) | cervical adenocarcinoma (human) | cervical squamous cell carcinoma (human) | endometrial adenocarcinoma (human) |
|---|---|---|---|---|---|---|---|---|---|---|---|---|---|---|---|---|---|---|---|---|---|---|---|---|---|---|---|
| B cell non Hodgkin lymphoma (human) | | | | | | | | | | | | | | | | | | | | | | | | | | | |
| chronic lymphocytic leukemia (human) | 0.628 | | | | | | | | | | | | | | | | | | | | | | | | | | |
| lymphoma (human) | 0.042 | 0.591 | | | | | | | | | | | | | | | | | | | | | | | | | |
| glioblastoma multiforme (human) | 0.746 | 0.954 | 0.801 | | | | | | | | | | | | | | | | | | | | | | | | |
| glioma (human) | 0.845 | 0.959 | 0.874 | 0.073 | | | | | | | | | | | | | | | | | | | | | | | |
| breast adenocarcinoma (human) | 0.886 | 0.957 | 0.959 | 0.491 | 0.636 | | | | | | | | | | | | | | | | | | | | | | |
| invasive lobular carcinoma (human) | 0.858 | 0.921 | 0.930 | 0.575 | 0.667 | 0.034 | | | | | | | | | | | | | | | | | | | | | |
| esophageal adenocarcinoma (human) | 0.828 | 0.778 | 0.959 | 0.491 | 0.633 | 0.373 | 0.460 | | | | | | | | | | | | | | | | | | | | |
| chromophobe renal cell carcinoma (human) | 0.846 | 0.804 | 0.955 | 0.673 | 0.813 | 0.622 | 0.769 | 0.175 | | | | | | | | | | | | | | | | | | | |
| renal cell carcinoma (human) | 0.632 | 0.884 | 0.714 | 0.156 | 0.179 | 0.667 | 0.724 | 0.485 | 0.568 | | | | | | | | | | | | | | | | | | |
| colorectal adenocarcinoma (human) | 0.827 | 0.790 | 0.950 | 0.522 | 0.720 | 0.467 | 0.576 | 0.066 | 0.242 | 0.562 | | | | | | | | | | | | | | | | | |
| cholangiocarcinoma (human) | 0.834 | 0.814 | 0.967 | 0.417 | 0.500 | 0.576 | 0.663 | 0.266 | 0.328 | 0.346 | 0.293 | | | | | | | | | | | | | | | | |
| hepatocellular carcinoma (human) | 1.108 | 1.079 | 1.131 | 1.007 | 0.988 | 1.076 | 1.089 | 1.047 | 1.005 | 1.032 | 1.042 | 0.625 | | | | | | | | | | | | | | | |
| lung adenocarcinoma (human) | 0.550 | 0.648 | 0.687 | 0.406 | 0.568 | 0.309 | 0.351 | 0.169 | 0.415 | 0.491 | 0.242 | 0.392 | 1.054 | | | | | | | | | | | | | | |
| squamous cell lung carcinoma (human) | 0.755 | 0.897 | 0.845 | 0.437 | 0.615 | 0.152 | 0.232 | 0.235 | 0.496 | 0.546 | 0.341 | 0.502 | 1.081 | 0.169 | | | | | | | | | | | | | |
| head and neck squamous cell carcinoma (human) | 0.822 | 0.842 | 0.910 | 0.449 | 0.540 | 0.240 | 0.277 | 0.219 | 0.562 | 0.541 | 0.379 | 0.522 | 1.082 | 0.169 | 0.060 | | | | | | | | | | | | |
| ovarian adenocarcinoma (human) | 0.832 | 0.705 | 0.944 | 0.652 | 0.716 | 0.245 | 0.202 | 0.485 | 0.656 | 0.754 | 0.571 | 0.613 | 1.064 | 0.336 | 0.500 | 0.517 | | | | | | | | | | | |
| pancreatic adenocarcinoma (human) | 0.544 | 0.775 | 0.652 | 0.413 | 0.558 | 0.291 | 0.319 | 0.220 | 0.443 | 0.533 | 0.336 | 0.478 | 1.068 | 0.068 | 0.214 | 0.204 | 0.373 | | | | | | | | | | |
| prostate adenocarcinoma (human) | 0.824 | 0.718 | 0.946 | 0.443 | 0.612 | 0.260 | 0.371 | 0.133 | 0.245 | 0.533 | 0.238 | 0.324 | 1.029 | 0.126 | 0.175 | 0.200 | 0.365 | 0.175 | | | | | | | | | |
| sarcoma (human) | 0.797 | 0.909 | 0.871 | 0.224 | 0.449 | 0.388 | 0.554 | 0.352 | 0.478 | 0.416 | 0.335 | 0.422 | 1.028 | 0.322 | 0.288 | 0.381 | 0.602 | 0.383 | 0.211 | | | | | | | | |
| melanoma (human) | 0.812 | 0.908 | 0.886 | 0.111 | 0.218 | 0.451 | 0.596 | 0.363 | 0.447 | 0.183 | 0.383 | 0.255 | 1.002 | 0.400 | 0.381 | 0.443 | 0.669 | 0.453 | 0.291 | 0.152 | | | | | | | |
| gastric adenocarcinoma (human) | 0.666 | 0.789 | 0.781 | 0.311 | 0.473 | 0.478 | 0.566 | 0.124 | 0.327 | 0.386 | 0.073 | 0.309 | 1.049 | 0.196 | 0.350 | 0.372 | 0.547 | 0.268 | 0.298 | 0.297 | 0.309 | | | | | | |
| follicular thyroid carcinoma (human) | 0.541 | 0.533 | 0.670 | 0.521 | 0.642 | 0.555 | 0.623 | 0.287 | 0.315 | 0.460 | 0.399 | 0.411 | 1.046 | 0.198 | 0.363 | 0.365 | 0.446 | 0.327 | 0.187 | 0.336 | 0.447 | 0.366 | | | | | |
| bladder transitional cell carcinoma (human) | 1.002 | 0.970 | 1.057 | 0.849 | 0.906 | 0.335 | 0.325 | 0.430 | 0.727 | 0.871 | 0.608 | 0.741 | 1.049 | 0.442 | 0.212 | 0.166 | 0.721 | 0.419 | 0.429 | 0.755 | 0.784 | 0.710 | 0.684 | | | | |
| cervical adenocarcinoma (human) | 0.987 | 0.905 | 1.062 | 0.872 | 0.903 | 0.650 | 0.654 | 0.617 | 0.776 | 0.726 | 0.679 | 0.826 | 1.104 | 0.637 | 0.690 | 0.633 | 0.535 | 0.690 | 0.641 | 0.740 | 0.815 | 0.726 | 0.722 | 0.811 | | | |
| cervical squamous cell carcinoma (human) | 0.853 | 0.854 | 0.940 | 0.565 | 0.687 | 0.219 | 0.273 | 0.202 | 0.492 | 0.617 | 0.346 | 0.531 | 1.087 | 0.192 | 0.045 | **0.032** | 0.508 | 0.237 | 0.195 | 0.419 | 0.499 | 0.395 | 0.378 | 0.144 | 0.586 | | |
| endometrial adenocarcinoma (human) | 0.976 | 0.885 | 1.077 | 0.656 | 0.787 | 0.141 | 0.153 | 0.401 | 0.637 | 0.755 | 0.409 | 0.569 | 1.074 | 0.426 | 0.362 | 0.452 | 0.158 | 0.512 | 0.399 | 0.505 | 0.584 | 0.458 | 0.582 | 0.597 | 0.524 | 0.395 | |

Yellow fill + bold - the minimum for human and dog
Violet fill + bold - human/canine distances below the median for human/human
Green fill + bold - human/canine distances below the median for both human/human and canine/canine
Orange fill + bold - human/canine distances below the minimal canine/canine distance
Red fill + bold - human/canine distances below the minimal human/human distance

**Table S5: Data sources for the RNAseq analysis;** FFPE – formalin-fixed paraffin-embedded; SE - single-end sequencing; na – not applicable, ns - not specified; if the dataset contained more samples but only a subset passed the criteria or was available, the total number was put in brackets.

| Study ID | Cancer type | Short | # of dogs | Exception | Dataset access | REF |
|---|---|---|---|---|---|---|
| PMID: 29674676 | B-cell lymphoma | BCL | 12(16) | - | GSE112474 | [22] |
| PMID: 30684308 | T-cell lymphoma | TCL | 6 | - | GSE122347 | [23] |
| NCATS-COP01 | T-cell lymphoma | TCL | 12 | - | caninecommons.cancer.gov/ | - |
| PMID: 32049048 | Glioma | Glioma | 42(83) | - | PRJNA579792 | [1] |
| PMID: 31570656 | Hemangiosarcoma | HSA | 8(23) | - | PRJNA562916 | [24] |
| PMID: 24525151 | Hemangiosarcoma | HSA | 45(51) | Data not shared in the paper | PRJNA376380 | [25] |
| PMID: 30089113 | Invasive urothelial carcinoma | iUC | 29 | - | GSE110661 | [26] |
| PMID: 31413331 | Mammary tumors | Breast | 158 | - | GSE119810 | [27] |
| NCATS-COP01 | Oral melanoma | OM | 12 | - | caninecommons.cancer.gov/ | - |
| PMID: 29066513 | Osteosarcoma | OS | 31 | - | GSE87649 | [28] |
| NCATS-COP01 | Pulmonary neoplasm | Lung | 12 | - | caninecommons.cancer.gov/ | - |
| PMID: 31041834 | Ameloblastoma | AM | 11(12) | SE | PRJNA533473 | [29] |
| PMID: 32665562 | Insulinoma | INS | 6 | Ns RNA library | PRJNA574196 | [30] |
| PMID: 29073243 | Meningioma | MEN | 13 | SE + FFPE | GSE95048 | [31] |
| PMID: 33142242 | Oral squamous cell carcinoma | OSCC | 10 | FFPE | PRJEB34234 | [32] |
| PMID: 31519932 | Prostate cancer | Prostate | 11 | Total RNA library | GSE122916 | [33] |

**Table S6: In this study, we analyzed the transcript expression for established and emerging immune checkpoints (ICs).** ICs were classified by their impact on the immune reaction: inhibitory, stimulatory, or complex. Identity [%] and coverage [%] were calculated by BlastP alignment of canine and human proteins; conservation score was calculated as identity*coverage/100. Receptors and their canonical ligands were grouped and marked with the same shades of gray.

| | Gene | Protein name | Description | Canine Ensembl ID | Identity | Coverage | Conservation |
|---|---|---|---|---|---|---|---|
| Inhibitory ICs | *PDCD1* | PD-1 | Programmed cell death 1 | ENSCAFG00000013184 | 72 | 39 | 28 |
| | *PDCD1LG1* | PD-L1 | Programmed cell death 1 ligand 1 | ENSCAFG00000002120 | 76 | 100 | 76 |
| | *PDCD1LG2* | PD-L2 | Programmed cell death 1 ligand 2 | ENSCAFG00000002121 | 67 | 95 | 64 |
| | *HAVCR1* | TIM-3 | Hepatitis A virus cellular receptor 1 | ENSCAFG00000023455 | 40 | 59 | 24 |
| | *LGALS9* | GAL-9 | Galectin 9 | ENSCAFG00000018641 | 72 | 91 | 66 |
| | *LAG3* | LAG-3 | Lymphocyte Activation Gene 3 | ENSCAFG00000014675 | 79 | 95 | 75 |
| | *FGL1* | FGL-1 | Fibrinogen like 1 | ENSCAFG00000032313 | 88 | 100 | 88 |
| | *CTLA4* | CTLA-4 | Cytotoxic T-lymphocyte associated protein 4 | ENSCAFG00000012876 | 87 | 100 | 87 |
| | *CD28* | CD28 | CD28 molecule | ENSCAFG00000012872 | 80 | 82 | 66 |
| | *CD80* | CD80 | CD80 molecule | ENSCAFG00000010997 | 54 | 100 | 54 |
| | *CD86* | CD86 | CD86 molecule | ENSCAFG00000011751 | 62 | 83 | 51 |
| | *TIGIT* | TIGIT | T cell immunoreceptor with Ig and ITIM domains | ENSCAFG00000010817 | 67 | 100 | 67 |
| | *NECTIN4* | NECTIN4 | Nectin Cell Adhesion Molecule 4 | ENSCAFG00000012722 | 94 | 100 | 94 |
| | *PVR* | CD155 | Poliovirus receptor | ENSCAFG00000004666 | 57 | 96 | 55 |
| | *SIRPA* | SIRPA | Signal-regulatory protein alpha | ENSCAFG00000025524 | 74 | 100 | 74 |
| | *CD47* | CD47 | Integrin associated protein | ENSCAFG00000009887 | 66 | 91 | 60 |
| | *PVRIG* | PVRIG | Poliovirus receptor-related immunoglobulin domain-containing | ENSCAFG00000014498 | 58 | 98 | 57 |
| | *VSIR* | VISTA | V-domain Ig suppressor of T-cell activation | ENSCAFG00000014354 | 82 | 100 | 82 |
| | *VTCN1* | B7-H4 | V-set domain-containing T-cell activation inhibitor 1 | ENSCAFG00000009858 | 94 | 100 | 94 |
| | *ADORA2A* | A2AR | Adenosine A2A receptor | ENSCAFG00000013828 | 93 | 100 | 93 |
| | *CYBB* | NOX2 | NADPH oxidase 2 | ENSCAFG00000013933 | 93 | 100 | 93 |
| Stimulatory ICs | *ICOS* | ICOS | Inducible T cell costimulator | ENSCAFG00000012880 | 70 | 99 | 69 |
| | *ICOSLG* | ICOSLG | Inducible T cell costimulator ligand | ENSCAFG00000010718 | 62 | 50 | 31 |
| | *TNFRSF4* | OX40 | TNF receptor superfamily member 4 | ENSCAFG00000019328 | 64 | 64 | 41 |
| | *TNFSF4* | OX40L | TNF superfamily member 4 | ENSCAFG00000014587 | 68 | 99 | 67 |
| | *TNFRSF9* | 4-1BB | TNF receptor superfamily member 9 | ENSCAFG00000019673 | 78 | 100 | 78 |
| | *TNFSF9* | 4-1BBL | TNF superfamily member 9 | ENSCAFG00000030400 | 66 | 58 | 38 |
| | *CD40* | CD40 | CD40 molecule | ENSCAFG00000009994 | 68 | 90 | 61 |
| | *CD40LG* | CD40L | CD40 ligand | ENSCAFG00000018945 | 83 | 100 | 83 |
| | *CD27* | CD27 | CD27 molecule | ENSCAFG00000015149 | 72 | 98 | 71 |
| | *CD70* | CD70 | CD70 molecule | ENSCAFG00000030308 | 69 | 100 | 69 |
| | *CD226* | CD226 | CD226 molecule | ENSCAFG00000000038 | 65 | 100 | 65 |
| Complex ICs | *TNFRSF18* | GITR | Glucocorticoid-induced TNFR-related protein | ENSCAFG00000019329 | 65 | 86 | 56 |
| | *TNFSF18* | GITRL | GITR ligand | ENSCAFG00000014590 | 72 | 100 | 72 |
| | *TNFRSF14* | HVEM | Herpesvirus entry mediator | ENSCAFG00000019422 | 58 | 80 | 46 |
| | *TNFSF14* | LIGHT | Homologous to lymphotoxin | ENSCAFG00000018627 | 82 | 100 | 82 |
| | *CD160* | CD160 | CD160 molecule | ENSCAFG00000055618 | 73 | 64 | 47 |
| | *BTLA* | BTLA | B- and T-lymphocyte attenuator | ENSCAFG00000010480 | 62 | 100 | 62 |
| | *KLRC1* | NKG2A | Killer cell lectin like receptor C1 | ENSCAFG00000028587 | 55 | 69 | 38 |
| | *CD276* | B7-H3 | CD276 molecule | ENSCAFG00000017820 | 94 | 60 | 56 |
| | *CD96* | TACTILE | T cell activation, increased late expression | ENSCAFG00000010358 | 69 | 100 | 69 |
| IC-like | *SLAMF7* | SLAMF7 | SLAM Family Member 7 | ENSCAFG00000012598 | 60 | 98 | 59 |
| | *ARG1* | Arginase 1 | Arginase 1 | ENSCAFG00000000386 | 92 | 100 | 92 |
| | *IDO1* | IDO | indoleamine 2,3-dioxygenase 1 | ENSCAFG00000005750 | 68 | 97 | 66 |

**Table S7: R configuration:** versions of the packages directly used in the projects as well as their dependencies

| package | version | package | version | package | version |
|---|---|---|---|---|---|
| askpass | 1.1 | grDevices | 4.2.3 | RColorBrewer | 1.1.3 |
| backports | 1.4.1 | grid | 4.2.3 | readr | 2.1.4 |
| base64enc | 0.1.3 | gtable | 0.3.2 | readxl | 1.4.2 |
| bit | 4.0.5 | haven | 2.5.2 | rematch | 1.0.1 |
| bit64 | 4.0.5 | highr | 0.10 | rematch2 | 2.1.2 |
| blob | 1.2.4 | hms | 1.1.2 | reprex | 2.0.2 |
| broom | 1.0.4 | htmltools | 0.5.4 | rlang | 1.1.0 |
| bslib | 0.4.2 | httr | 1.4.5 | rmarkdown | 2.20 |
| cachem | 1.0.7 | ids | 1.0.1 | rstudioapi | 0.14 |
| callr | 3.7.3 | isoband | 0.2.7 | rvest | 1.0.3 |
| cellranger | 1.1.0 | jquerylib | 0.1.4 | sass | 0.4.5 |
| cli | 3.6.0 | jsonlite | 1.8.4 | scales | 1.2.1 |
| clipr | 0.8.0 | knitr | 1.42 | selectr | 0.4.2 |
| colorspace | 2.1.0 | labeling | 0.4.2 | splines | 4.2.3 |
| conflicted | 1.2.0 | lattice | 0.20.45 | stats | 4.2.3 |
| cpp11 | 0.4.3 | lifecycle | 1.0.3 | stringi | 1.7.12 |
| crayon | 1.5.2 | lubridate | 1.9.2 | stringr | 1.5.0 |
| curl | 5.0.0 | magrittr | 2.0.3 | sys | 3.4.1 |
| data.table | 1.14.8 | MASS | 7.3.58.3 | systemfonts | 1.0.4 |
| DBI | 1.1.3 | Matrix | 1.5.3 | textshaping | 0.3.6 |
| dbplyr | 2.3.1 | memoise | 2.0.1 | tibble | 3.2.1 |
| digest | 0.6.31 | methods | 4.2.3 | tidyr | 1.3.0 |
| dplyr | 1.1.0 | mgcv | 1.8.42 | tidyselect | 1.2.0 |
| dtplyr | 1.3.0 | mime | 0.12 | tidyverse | 2.0.0 |
| ellipsis | 0.3.2 | modelr | 0.1.10 | timechange | 0.2.0 |
| evaluate | 0.20 | munsell | 0.5.0 | tinytex | 0.44 |
| fansi | 1.0.4 | nlme | 3.1.162 | tools | 4.2.3 |
| farver | 2.1.1 | openssl | 2.0.6 | tzdb | 0.3.0 |
| fastmap | 1.1.1 | pillar | 1.8.1 | utf8 | 1.2.3 |
| forcats | 1.0.0 | pkgconfig | 2.0.3 | utils | 4.2.3 |
| fs | 1.6.1 | prettyunits | 1.1.1 | uuid | 1.1.0 |
| gargle | 1.3.0 | processx | 3.8.0 | vctrs | 0.6.0 |
| generics | 0.1.3 | progress | 1.2.2 | viridisLite | 0.4.1 |
| ggplot2 | 3.4.1 | ps | 1.7.2 | vroom | 1.6.1 |
| glue | 1.6.2 | purrr | 1.0.1 | withr | 2.5.0 |
| googledrive | 2.0.0 | R6 | 2.5.1 | xfun | 0.39 |
| googlesheets4 | 1.0.1 | ragg | 1.2.5 | xml2 | 1.3.3 |
| graphics | 4.2.3 | rappdirs | 0.3.3 | yaml | 2.3.7 |

#### Supplementary figures

**Figure S1: IC abundance plot with values normalized to mean expression of each gene, corresponding to Fig. 1A;** each value was divided by a mean of the abundances of the respective gene across all cancer types to obtain the visualized abundance score. This way the relative up- or down-regulation of each IC in between the cancer types can be inspected without the confusion caused by transcript abundances naturally characterized by different orders of magnitude depending on the IC; gray color - lack of information due to undetectable expression.

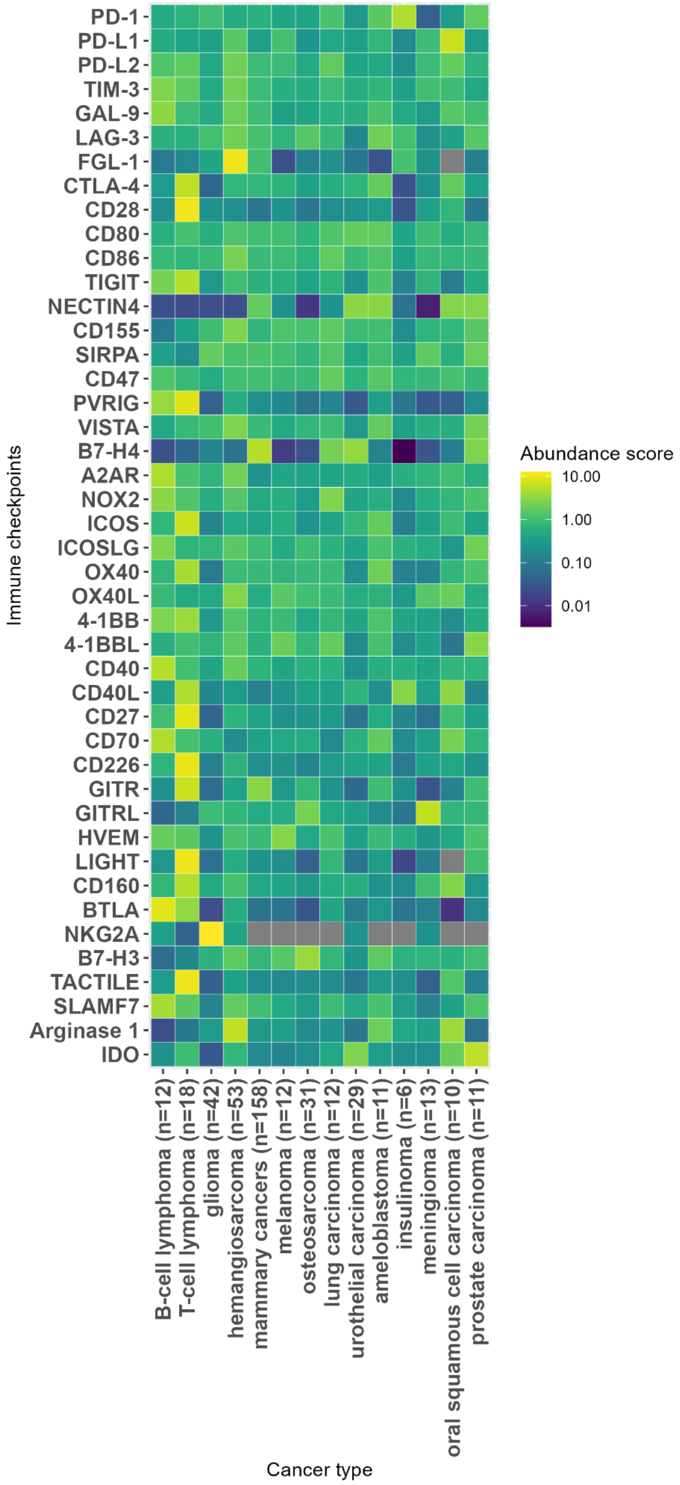

**Figure S2: Variance control plot corresponding to the median IC abundance plot (Fig. 1A):** Median Absolute Deviation divided by median (MAD/m); grey colour - lack of information due to undetectable expression.

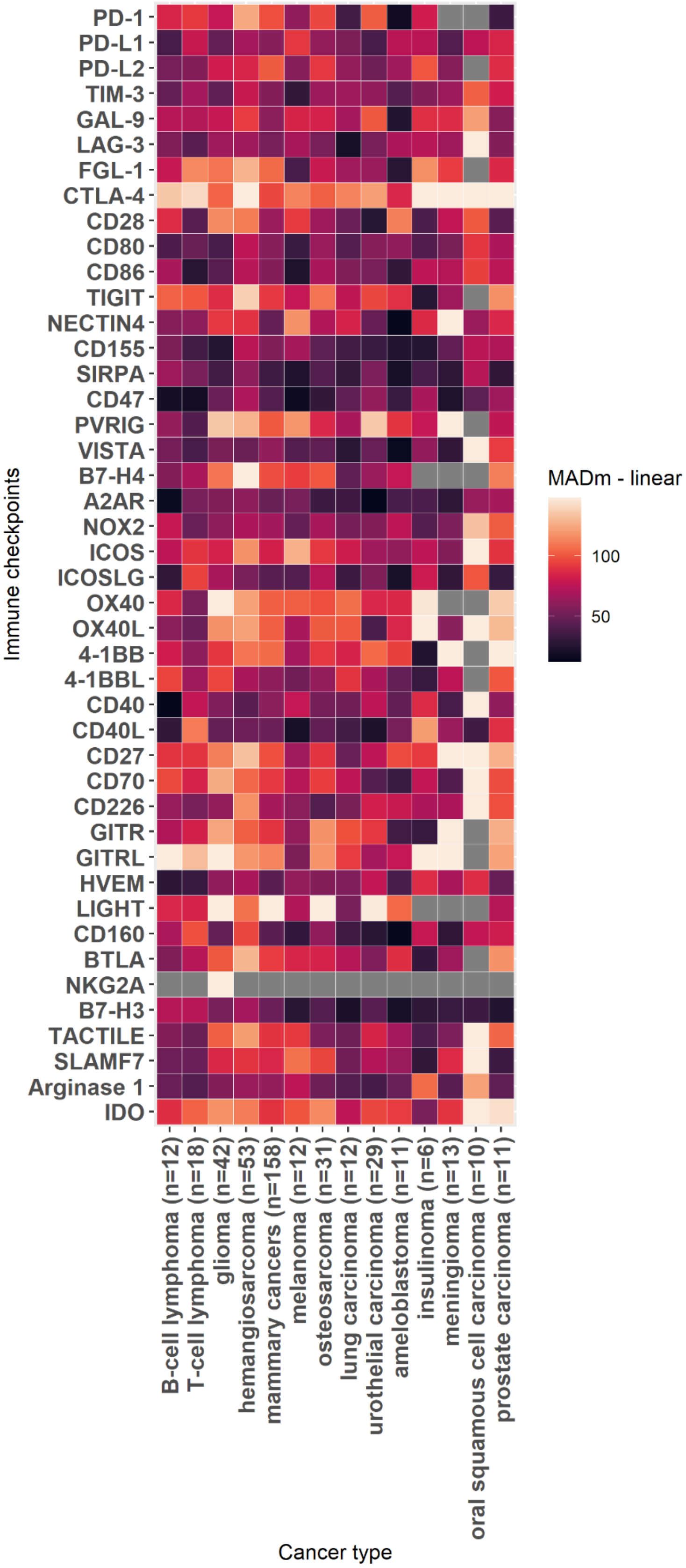

**Figure S3: Median TPM-quantified expression of ICs across human cancers based on EBI-obtained human data;** gray color - lack of information due to undetectable expression.

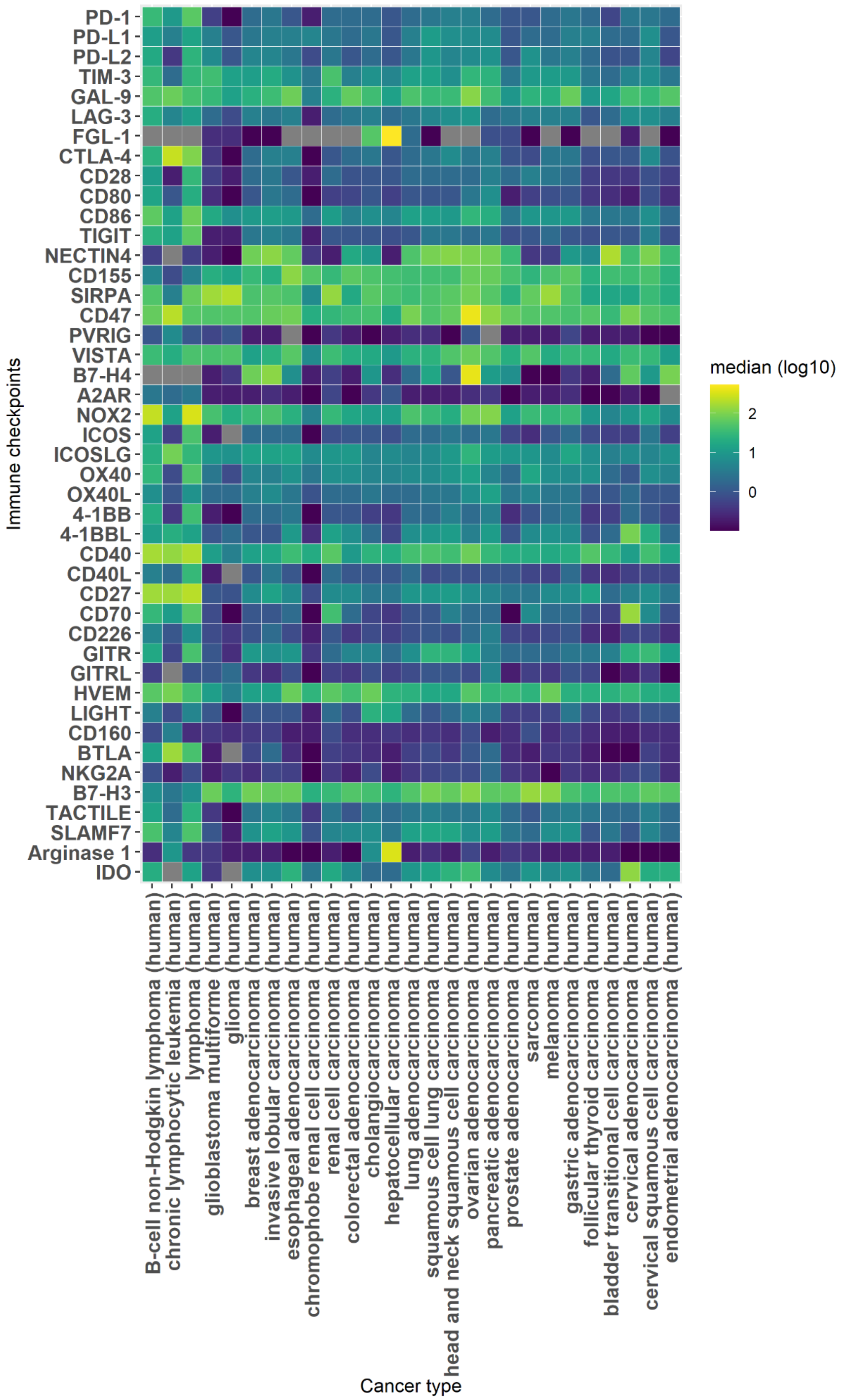

**Figure S4: Boxplots representing the distribution of IC expression across individual samples in each canine cancer type.**

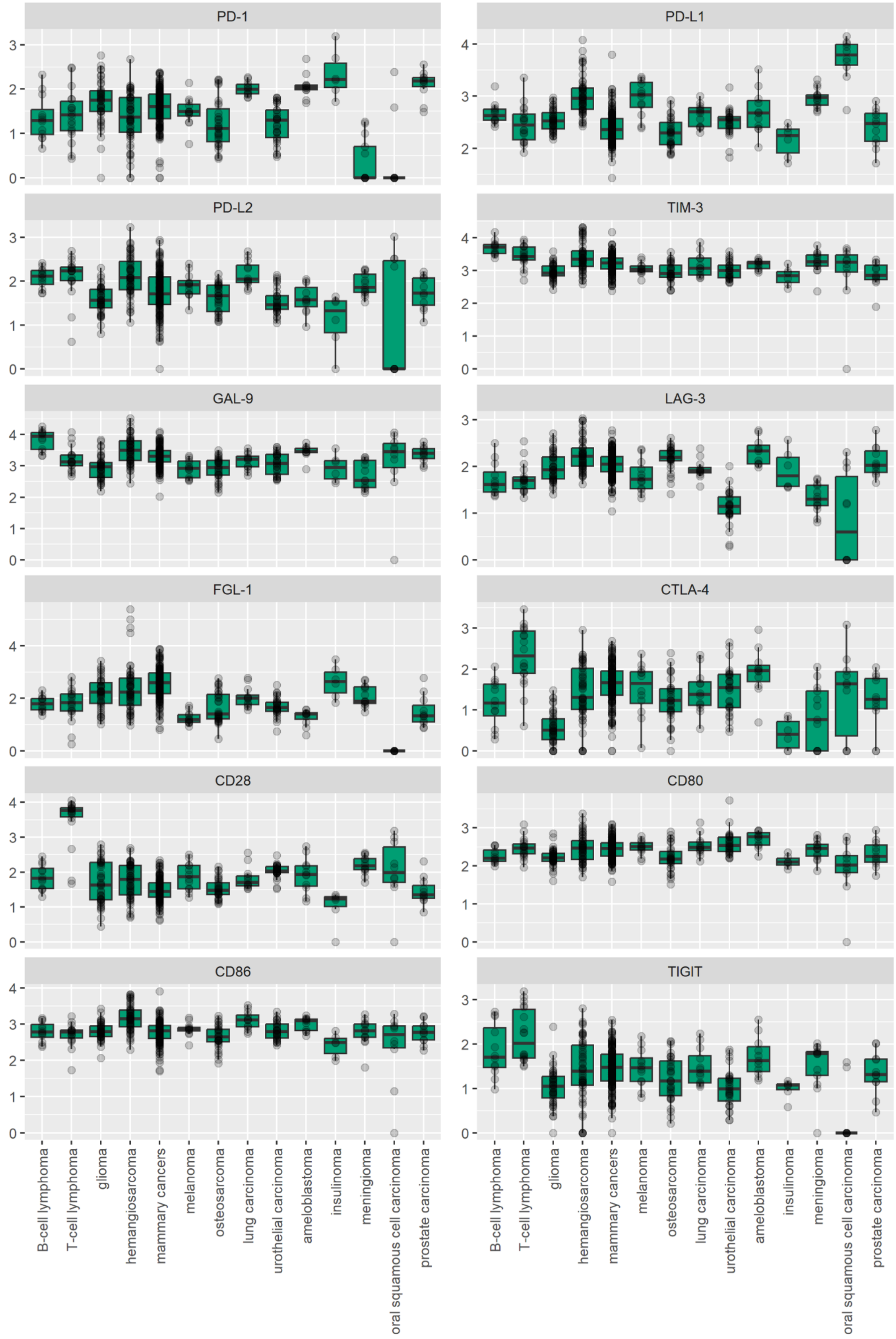

Fig. S4 - Continued.

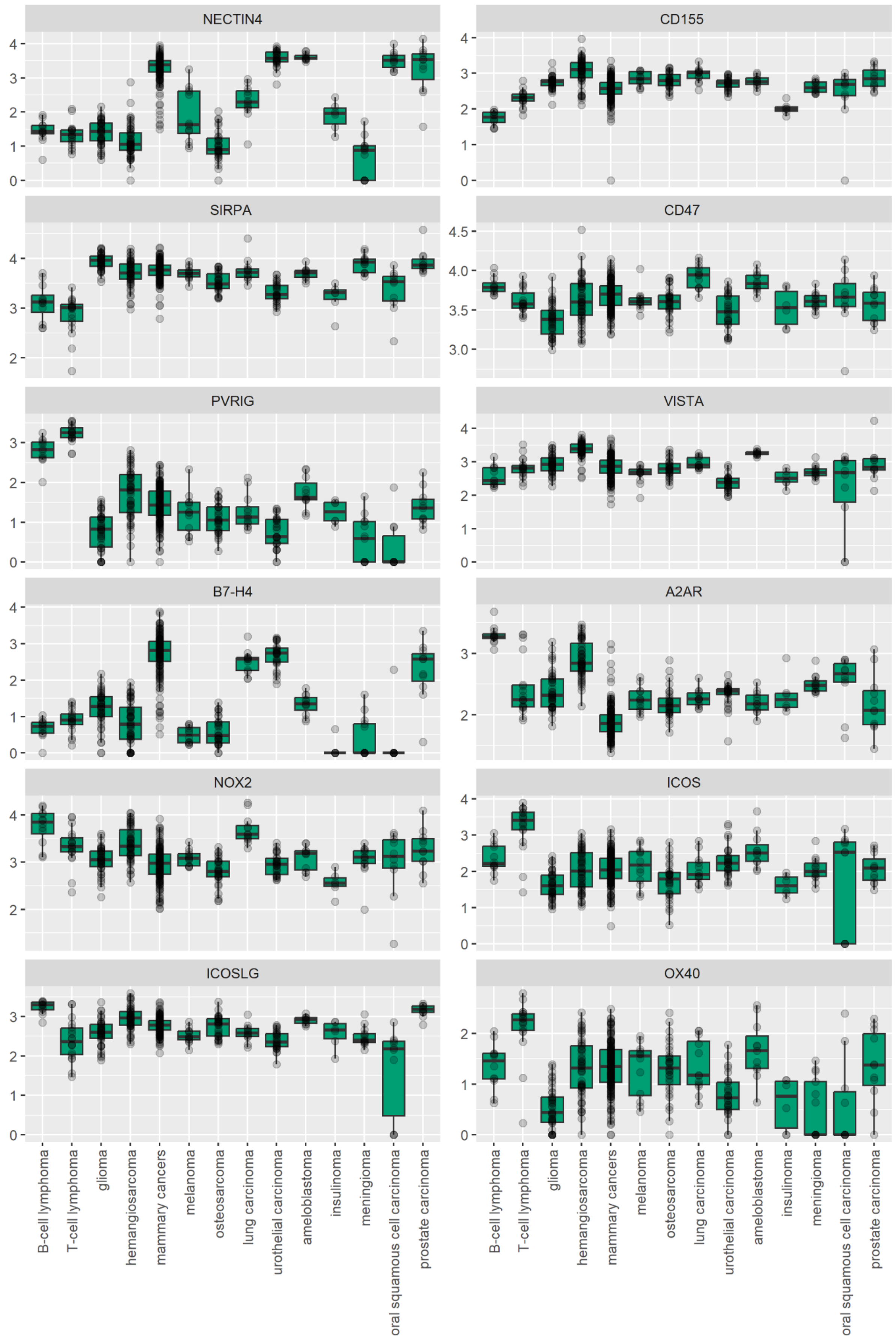

Fig. S4 - Continued.

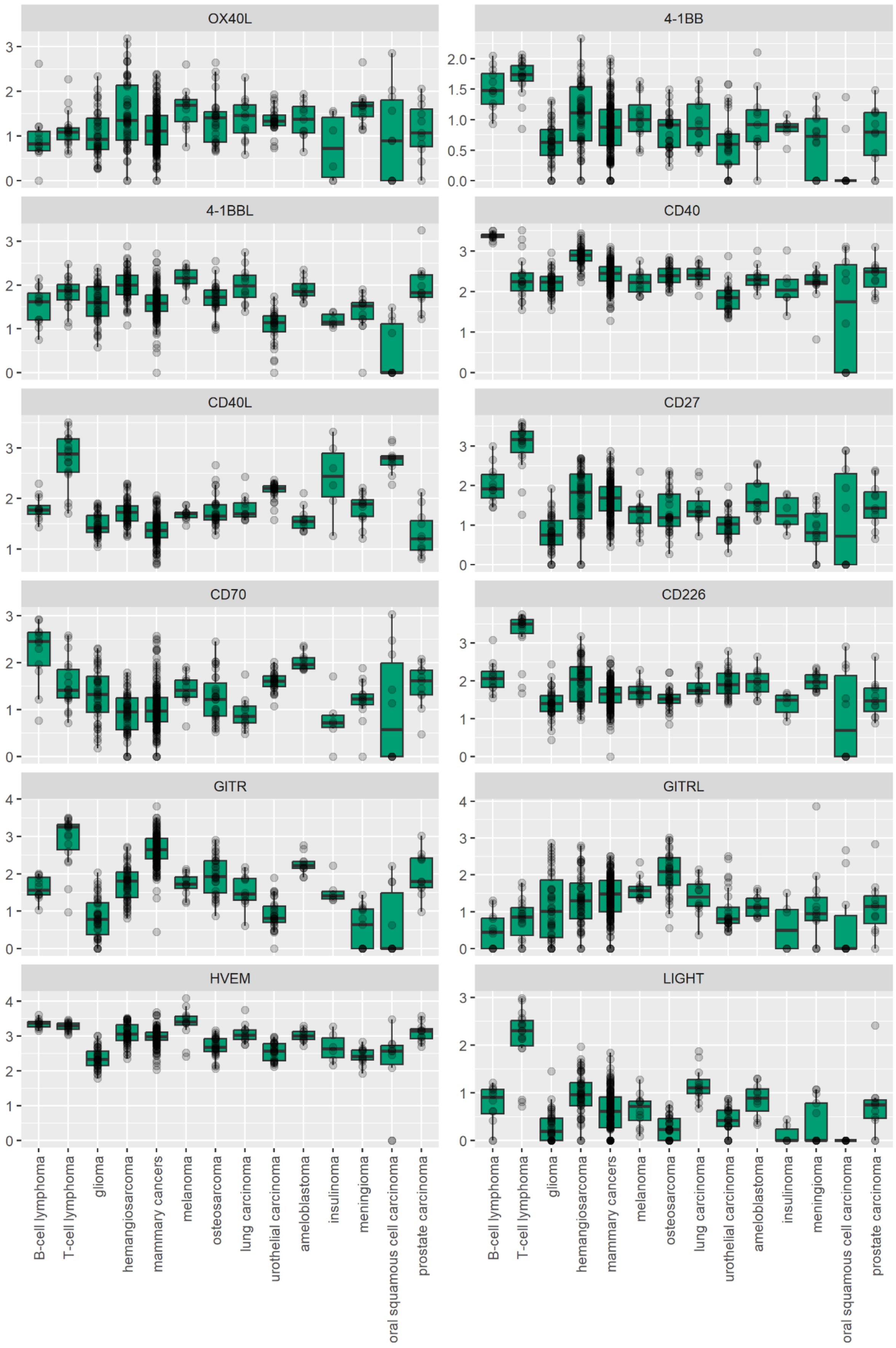

Fig. S4 - Continued.

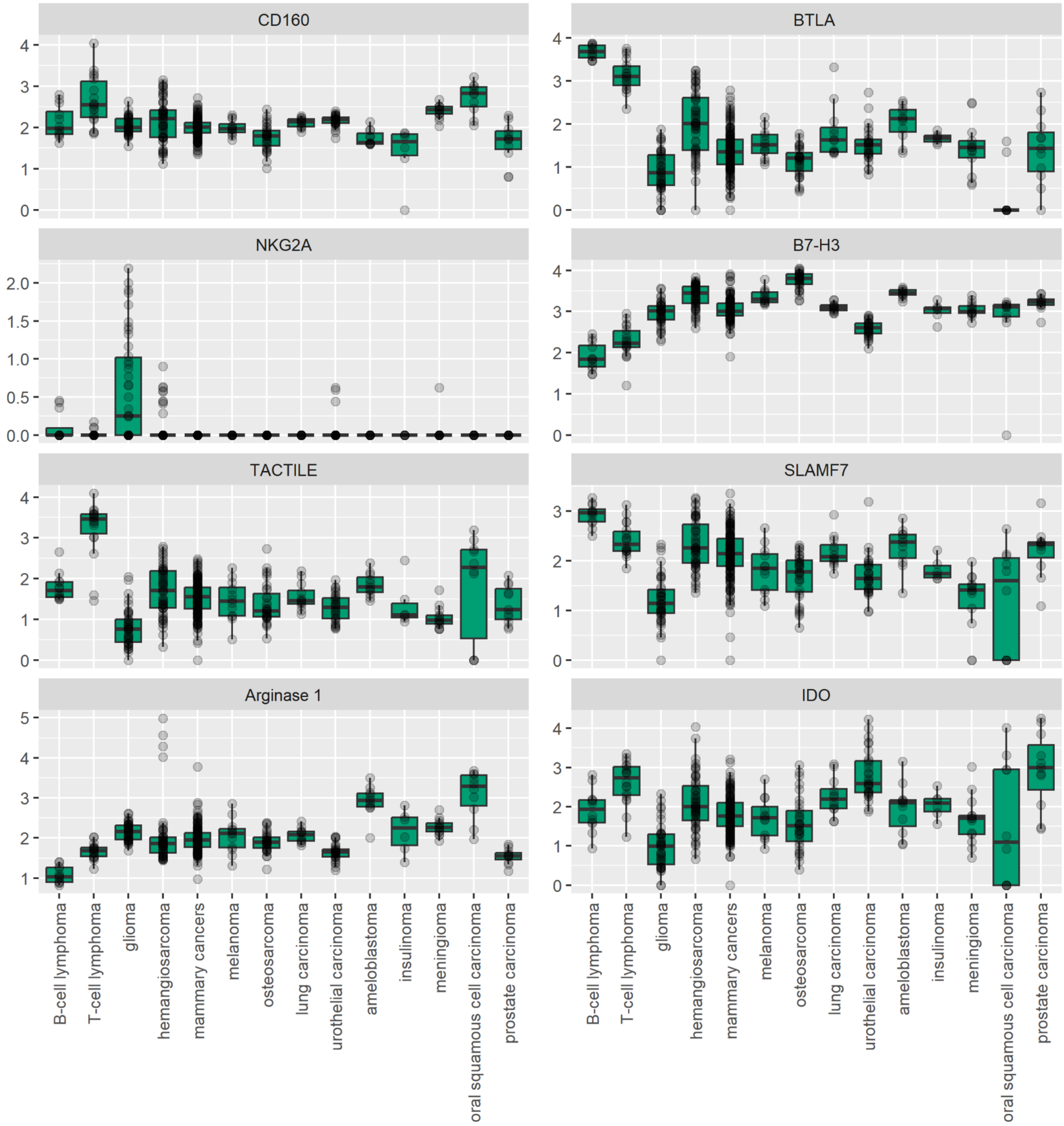

**Figure S5: Normalized expression of the markers of immune cell populations infiltrating tumors.**

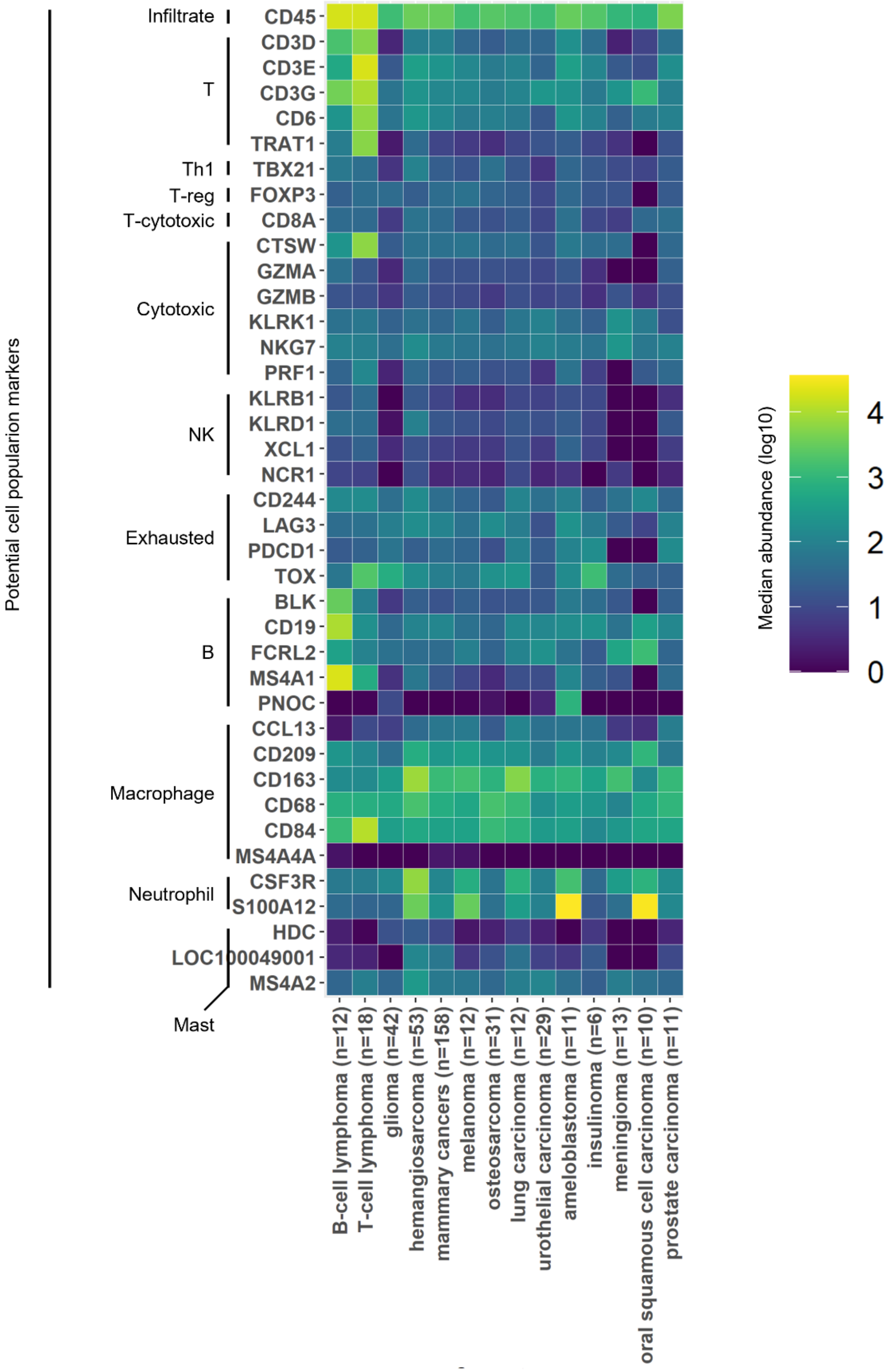

**Figure S6: UMAP representation of individual patient IC signatures across the human/canine cancers.**

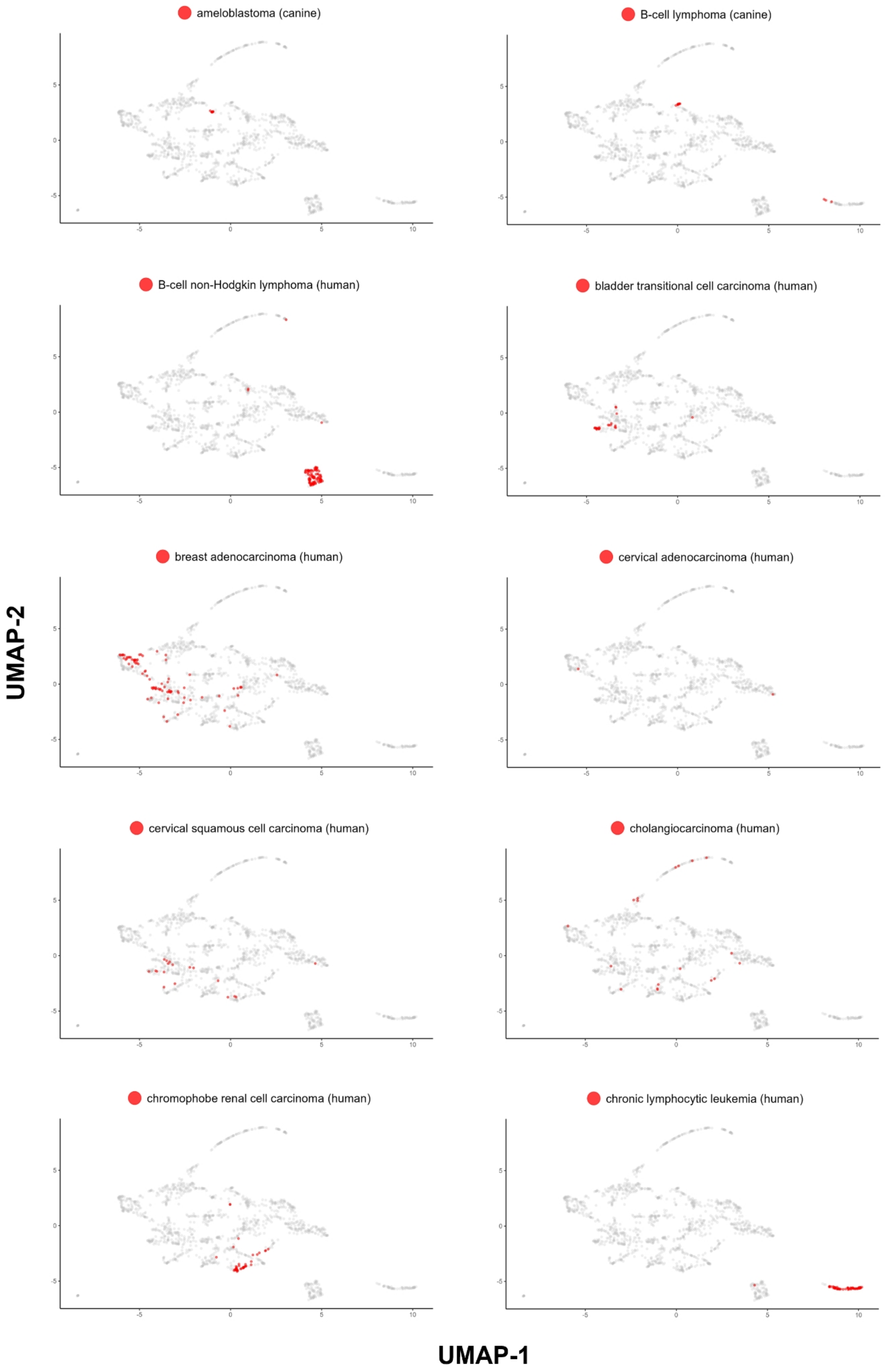

Fig. S6 - Continued.

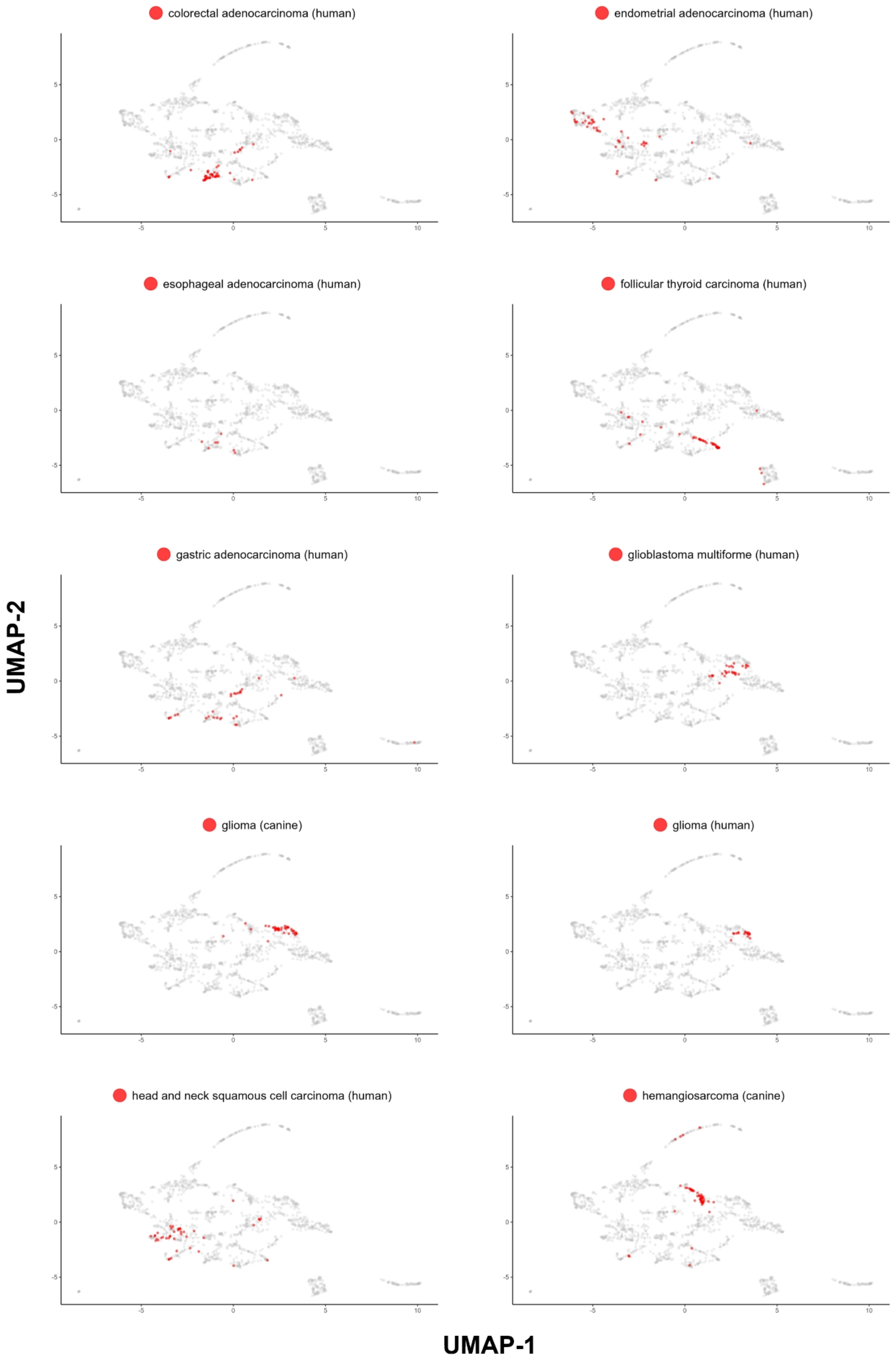

Fig. S6 - Continued.

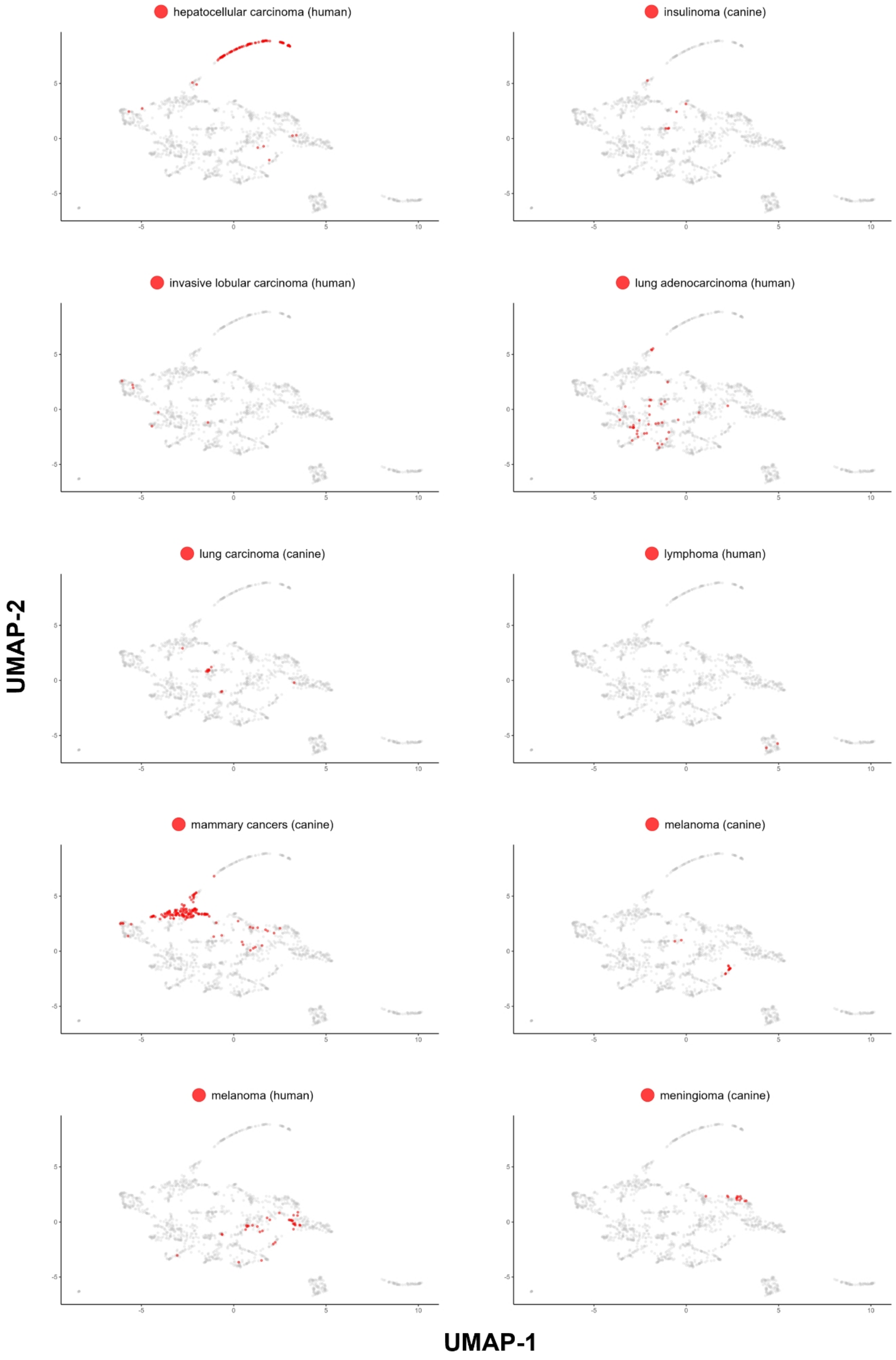

Fig. S6 - Continued.

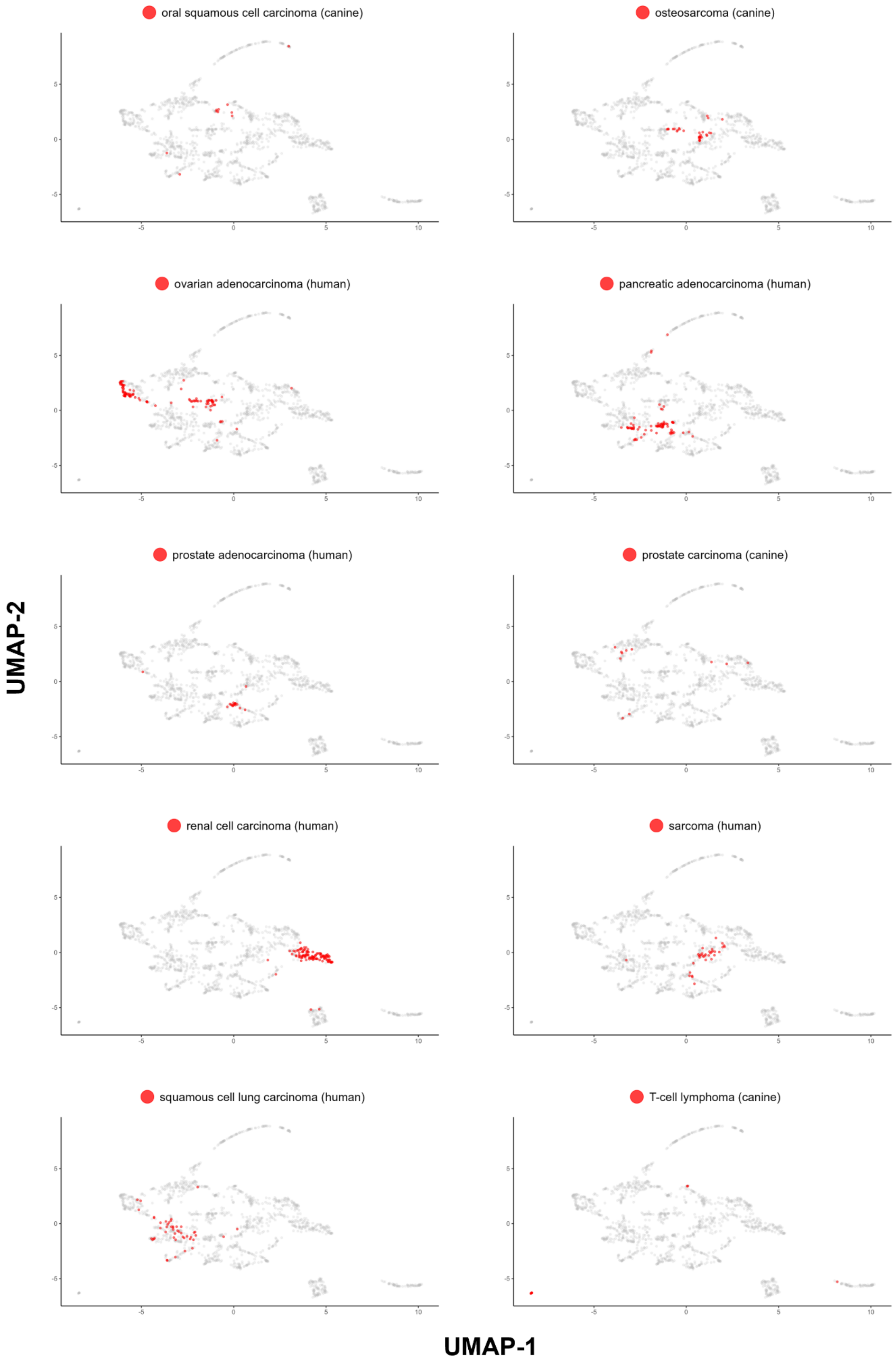

Fig. S6 - Continued.

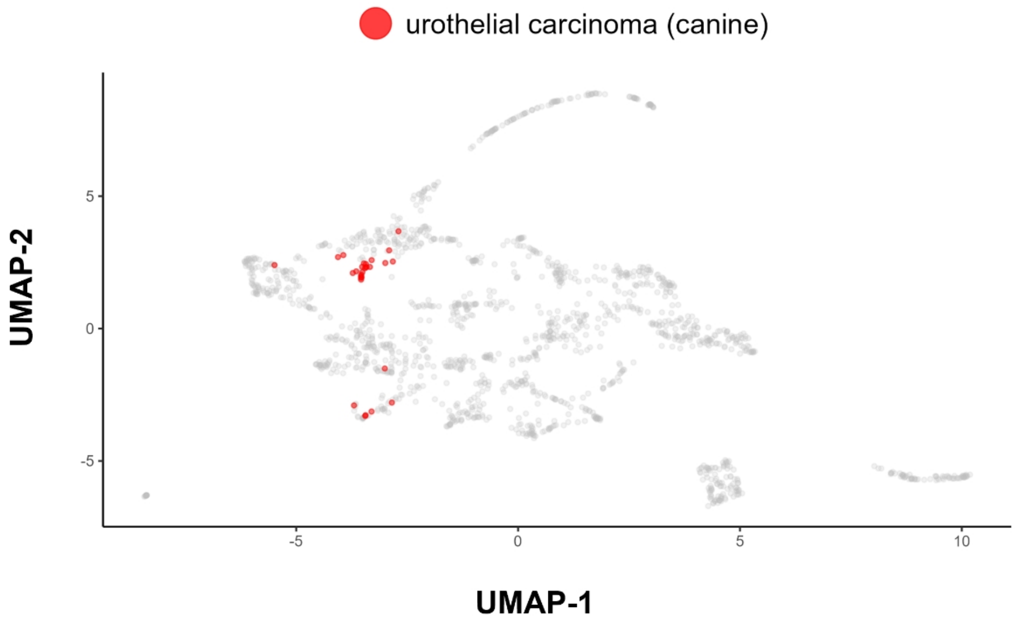


**Figure S7: A dendrogram of human and canine cancer types corresponding to Fig. 4F, but prepared by clustering the data by 4 randomly chosen IC genes only**; **blue** - human and **green** - canine cancers.

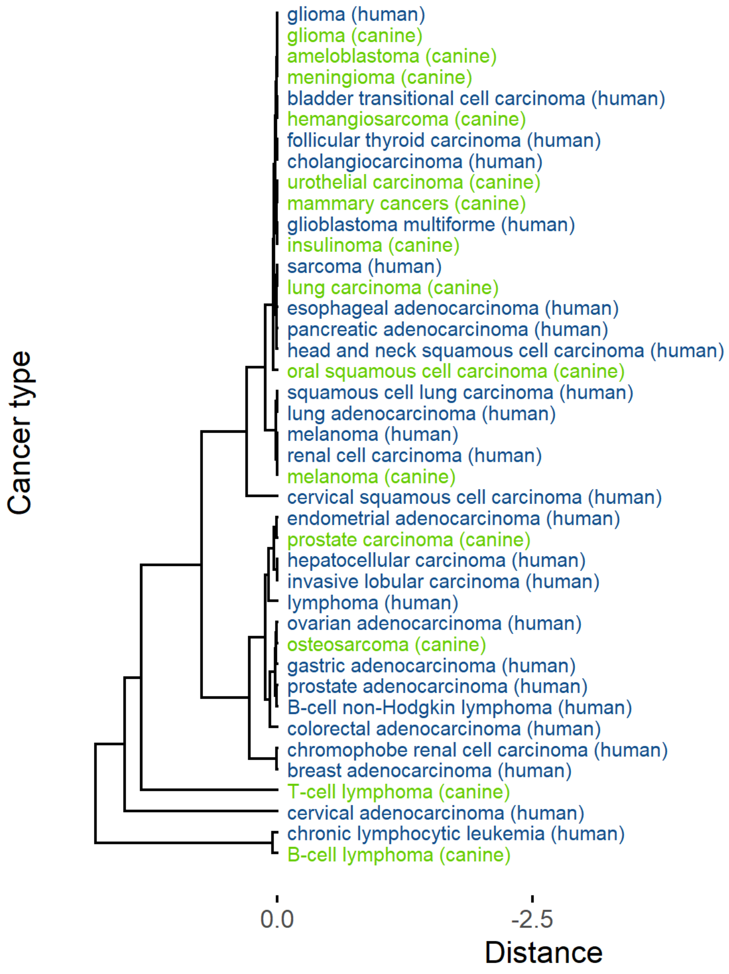
